# The amplitude of gammaherpesvirus lytic replication dictates adaptive immune activation: Potential implications for KSHV LANA in immune evasion

**DOI:** 10.64898/2026.02.05.703987

**Authors:** Steven J. Murdock, Shana M. Owens, Darby G. Oldenburg, J. Craig Forrest

## Abstract

Adaptive immune responses to primary Kaposi sarcoma-associated herpesvirus (KSHV) infection are poorly defined. To develop better small-animal models for understanding KSHV pathogenesis and immunity, we previously generated a chimeric virus in which the KSHV latency-associated nuclear antigen (kLANA), a conserved multifunctional protein critical for viral latency, was exchanged for the LANA homolog in murine gammaherpesvirus 68 (MHV68). Despite comparable levels of latent infection between WT and KLKI MHV68, kLANA directly repressed MHV68 lytic replication and reactivation. We therefore hypothesized that suppression of lytic replication by kLANA dampens adaptive immune responses. To test this, mice were infected with equivalent doses of either WT or KLKI MHV68 and adaptive immune responses were evaluated over time. B and T cell activation was starkly reduced following KLKI MHV68 infection, despite a potent virus-specific effector CD8+ T cell response against both viruses. These phenotypes were independent of inoculating dose, as high dose infection with KLKI MHV68 still showed reduced adaptive immune cell activation. In contrast, infection of *Ifnar1^-/-^*mice, which support enhanced KLKI MHV68 lytic replication, led to potent adaptive cellular and humoral immune activation by both WT and KLKI viruses, suggesting that the level of viral replication, and not simply amount of virus present, is a major driver of adaptive immunity during GHV infection. Collectively, these data support the hypothesis that kLANA-mediated suppression of lytic replication facilitates immune evasion by holding viral replication below a threshold for potent induction of adaptive immunity.

**IMPORTANCE:** KSHV is a gammaherpesvirus that establishes lifelong, chronic infections in humans and increases the risk of virus-associated cancers. Currently, there is little information on how primary KSHV infection influences adaptive immune development in healthy individuals. Rodent models, such as murine gammaherpesvirus 68 (MHV68), provide a valuable laboratory system for studying gammaherpesvirus pathogenesis *in vivo*. In this study, we report that infection with a previously characterized chimeric KSHV-MHV68 virus expressing KSHV LANA represses lytic viral replication and elicits weak antiviral adaptive immune responses following primary infection, despite efficient latency establishment. Using this chimeric MHV68 virus, we demonstrate that lytic viral amplification must breach a threshold to trigger a potent virus-specific adaptive immune response. We propose that KSHV, through LANA, evades detection by repressing lytic viral replication to remain “below the radar” of adaptive immune defenses during host colonization.

## INTRODUCTION

The gammaherpesvirus (GHV) subfamily of herpesviruses are large, enveloped viruses that contain a double-stranded DNA genome. GHVs exhibit a biphasic infection cycle, which includes a lytic and a latent stage of viral infection (1). The lytic, or productive, phase is characterized by viral genome replication and generation of viral progeny. Lytic replication allows GHVs to traverse host mucosal barriers for dissemination and transmission in a proinflammatory manner, ultimately resulting in cell death and virion production (2). After lytic infection is cleared, GHVs establish latency, a phase of infection in which the viral genome is maintained in the host-cell nucleus as an extrachromosomal, covalently closed dsDNA episome, and the production of viral progeny is ceased. The transition to latency allows the virus to evade immune surveillance and persist for the lifetime of the host, however specific stimuli can induce viral reactivation wherein the virus reinitiates lytic gene expression and restarts the productive cycle again. Latency is likely minimally immunostimulatory, as viral transcription is limited to a few genes that promote infected cell survival and viral genome maintenance within host cells.

Kaposi sarcoma-associated herpesvirus (KSHV, also known as human herpesvirus 8, HHV-8) is a human GHV that establishes latent infection in B lymphocytes and other cell types and is linked to lymphoproliferative malignances and cancer (3). For KSHV, seroprevalence varies based on geographical location, however, seropositivity ranges from 50-70% in endemic regions like sub-Saharan Africa and in people living with HIV infection (PLWH) (4). While GHV infections are not always linked to severe disease, KSHV-infected patients can develop primary effusion lymphoma, multicentric Castleman disease, the AIDS-defining endothelial cell malignancy, Kaposi sarcoma, and KSHV inflammatory cytokine syndrome (5).

It is virtually unknown how adaptive immune responses develop during primary KSHV infection in humans, and this is likely because primary KSHV infection is asymptomatic with no signs of clinical disease. Studies conducted in KSHV endemic populations indicate that healthy individuals develop relatively weak immune responses to KSHV infection. Antibody responses in individuals with low or no detectable HIV viral loads and no previous history of KSHV-associated diseases are weakly reactive to most of the KSHV proteome with few immunodominant targets (6). Furthermore, in KSHV endemic areas such as rural Uganda, the abundance of KSHV-specific CD4+ and CD8+ T cells is low in infected individuals and target only a few immunodominant peptides (7). These findings suggest that KSHV is well controlled in healthy individuals and may have evolved to establish persistent infection in humans without eliciting potent adaptive immune responses. While several immunomodulatory proteins are encoded by KSHV, which of these viral gene products help KSHV avoid potently stimulating adaptive immunity is not known, especially *in vivo*.

Since chronic GHV infection can lead to cancer, it is critical that we understand the viral factors that promote latent infection and immune evasion. For KSHV, the latency-associated nuclear antigen (LANA) is encoded by the *ORF73* gene and is essential for latency establishment and viral episome persistence by tethering the viral genome to host chromatin (8, 9). In addition to maintaining the viral genome in infected cells, KSHV LANA (kLANA) suppresses lytic replication by repressing the lytic switch protein replication and transcription activator (RTA) through modifications to viral chromatin during latency (10). Indeed, during *de novo* KSHV infection of SLK cells, kLANA recruits polycomb repressor complex 2 (PRC2) to the viral genome to silence lytic promoters, including the promoter for the RTA-encoding *ORF50* gene (11), suggesting that inhibiting lytic replication is an important event in primary KSHV infection and latency establishment.

Although GHVs exhibit a strict host restriction, several conserved proteins, including LANA, are functionally interchangeable between KSHV and murine gammaherpesvirus 68 (MHV68), a genetically related rodent virus frequently used to understand GHV pathogenesis *in vivo*. Indeed, using an MHV68 recombinant in which the MHV68 LANA (mLANA)-encoding *ORF73* gene was replaced with the kLANA-encoding *ORF73* gene, we and others demonstrated that mLANA and kLANA are functionally interchangeable to enable latency establishment and long-term maintenance following MHV68 infection of mice (12, 13). Despite efficient latency establishment by the kLANA knock-in (KLKI) MHV68, we also observed that kLANA, but not mLANA, repressed the MHV68 *ORF50* promoter and limited RTA expression and subsequent lytic replication and reactivation (12). This finding revealed that the capacity of kLANA to suppress productive viral replication was transferrable to MHV68. We also found that KLKI MHV68 did not cause splenomegaly, an infectious-mononucleosis (IM)-like pathology that is dependent on MHV68-driven CD4+ T cell activation (14).

In addition to promoting an IM-like syndrome, MHV68 generally elicits a potent virus-specific adaptive immune response that is characterized by widespread B and T cell activation targeting multiple viral antigens (15–17). In contrast to WT MHV68, replication-defective viruses like RTA-null (50.Stop) MHV68 minimally elicit adaptive immunity, requiring multiple high-dose inoculations to promote virus-specific immunity (18, 19). Similarly, blockade of MHV68 replication with antiviral drugs also prevents virus-specific immunity (20). Together these findings suggest the hypothesis that lytic GHV replication, and perhaps associated inflammation, is critical for stimulating virus-specific adaptive immunity. Given that kLANA suppresses lytic viral replication and immunity to primary KSHV infection is weak, we hypothesized that kLANA-mediated suppression of lytic replication is immunoevasive and serves to limit GHV-specific immune responses. By integrating MHV68 immunology with the KLKI MHV68 model, we sought to define how the amplitude of lytic replication, as dictated by kLANA, influences adaptive immune activation during MHV68 primary infection and latency establishment.

## RESULTS

### kLANA-mediated repression of *mORF50* restricts lytic antigen accumulation during MHV68 lytic replication

We previously demonstrated that kLANA, but not mLANA, has the capacity to repress MHV68 *ORF50* (mRTA) transcription. For a chimeric MHV68 virus in which mLANA was replaced with kLANA, this function correlated with suppressed MHV68 lytic replication, a phenotype that was overcome by providing mRTA *in trans* (12). To more broadly define how kLANA impacted MHV68 viral gene transcription, we performed quantitative RT-PCR (qRT-PCR) comparing WT, KLKI, and mLANA-deficient 73.Stop MHV68 transcription over time during lytic infection (21). Although LANA-null MHV68 is attenuated in murine fibroblasts following low-MOI infections, mLANA-null virus retains its capacity to progressively express viral gene products in a time-dependent manner (22, 23). Both WT and 73.Stop MHV68 exhibited the expected progression through the lytic transcription cascade, with viral transcripts increasing in abundance over time **(Fig. 1A)**. In contrast, KLKI MHV68 infection was characterized by low-level lytic transcription that did not appear to amplify as infection progressed. Immunoblot analyses of infected cell lysates supported this observation, as lytic viral antigens detected with MHV68 immune sera were less abundant in cells infected with KLKI MHV68 compared to WT and 73.Stop MHV68 **(Fig. 1B).** KLKI MHV68 also exhibited a small plaque phenotype compared to WT or 73.Stop infections **(Fig. 1C)**. Consistent with mRTA provided *in trans* rescuing KLKI MHV68 replication (12), infection of cells stably expressing mRTA equilibrated KLKI MHV68 transcription to levels approximating those of WT and 73.Stop MHV68 and enhanced the production of lytic viral antigens (**Fig. S1A-B**). These data indicate that expression of kLANA in place of mLANA correlates with suppressed viral transcription and decreased production of lytic antigens during productive MHV68 replication. In light of our prior work (12) and enhanced viral gene transcription and translation occurring in mRTA-expressing cells, this finding is consistent with kLANA mediating suppression of *ORF50* transcription.

**Figure 1.**
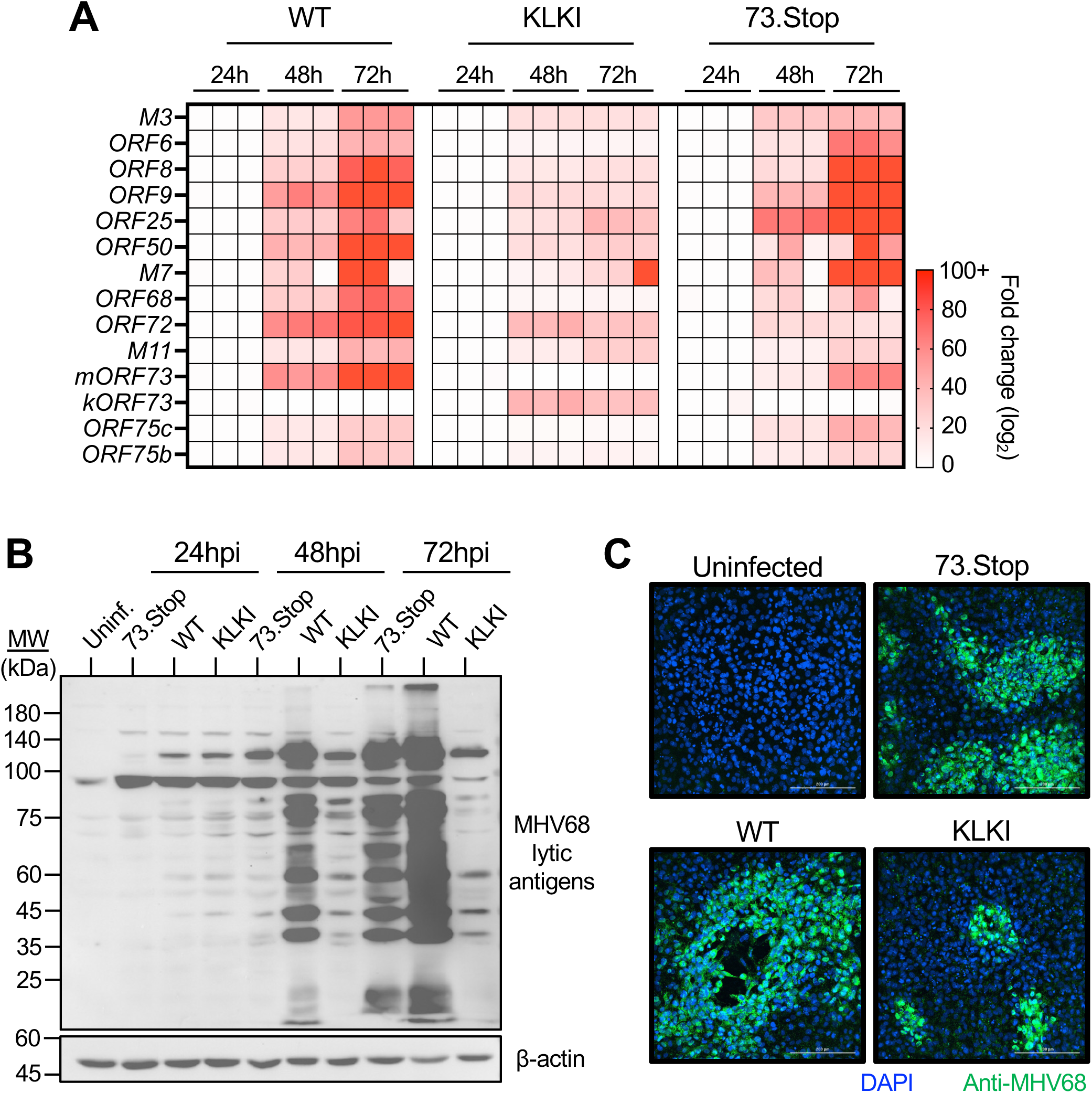
Expression of kLANA correlates with reduced lytic antigen accumulation during MHV68 infection. 3T12 fibroblasts were infected with the indicated viruses at an MOI of 0.1 PFU/cell. (A) RNA was collected at the indicated times postinfection. Viral transcripts were quantified by qRT-PCR relative to *B2m* using the ΔΔCT method and normalized to transcript abundance at 24hpi. (B) Lysates were collected at the indicated times postinfection and immunoblot analyses were performed using antibodies that recognize the indicated proteins. (C) Cells were fixed 72h postinfection and stained for indirect immunofluorescence analysis using antibodies that recognize the indicated proteins. DNA was stained with DAPI. Scale bar – 200uM.

### KLKI MHV68 infection minimally stimulates B and T cell activation and immunity

Similar to Epstein-Barr virus (EBV) infection of humans, infection of mice with WT MHV68 causes splenomegaly, an immune-driven pathology characterized by immune cell recruitment and proliferation resulting in an increase in spleen mass and cellularity (14). In contrast, KSHV is not known to cause splenomegaly during primary infection. We previously demonstrated that although KLKI MHV68 disseminated systemically and established latency in the spleen at levels equivalent to WT virus, KLKI MHV68 did not cause splenomegaly and only minimally reactivated from latency when compared to WT virus infections (12), phenotypes we confirmed in support of this study **(Fig. 4A,F and Fig. S2)**.

Since splenomegaly is an immune mediated pathology and cells infected with KLKI MHV68 generate fewer lytic viral antigens over time, we hypothesized that the presence of kLANA and the related suppression of lytic gene expression would reduce immune stimulation by KLKI MHV68 relative to WT virus. To test this hypothesis, we compared B and T cell populations in the spleen on days 16-18 after intranasal infection with 1000 PFU of either WT or KLKI MHV68. This dose of virus and timepoint typically correlates with potent immune activation after WT MHV68 infection. Flow cytometry analyses of total B cells (CD19^+^ B220^+^), T cells (CD3^+^), and CD4+ or CD8+ T cells (CD3^+^ CD4^+^ or CD8^+^) in spleens indicated no major differences in the total number of general lymphocyte populations between infections **(Fig. S3)**. When germinal center (GC) B cells (CD19^+^ B220^+^ GL7^+^ CD38^lo^) and T follicular helper cells (Tfh, CD4^+^ CXCR5^+^ PD-1^+^) were analyzed, expansion of both cell types was potently stimulated by WT virus, whereas neither cell type was significantly increased relative to naïve mice following KLKI MHV68 infection **(Fig. 2A-B)**. Plasmablast (CD138^+^ B220^lo^) expansion also was lower following KLKI MHV68 infection, however the reduction was not significant compared to WT virus **(Fig. 2C)**. Together, these data suggest that KLKI MHV68 infection establishes latency without triggering potent B cell activation and GC B cell expansion.

**Figure 2.**
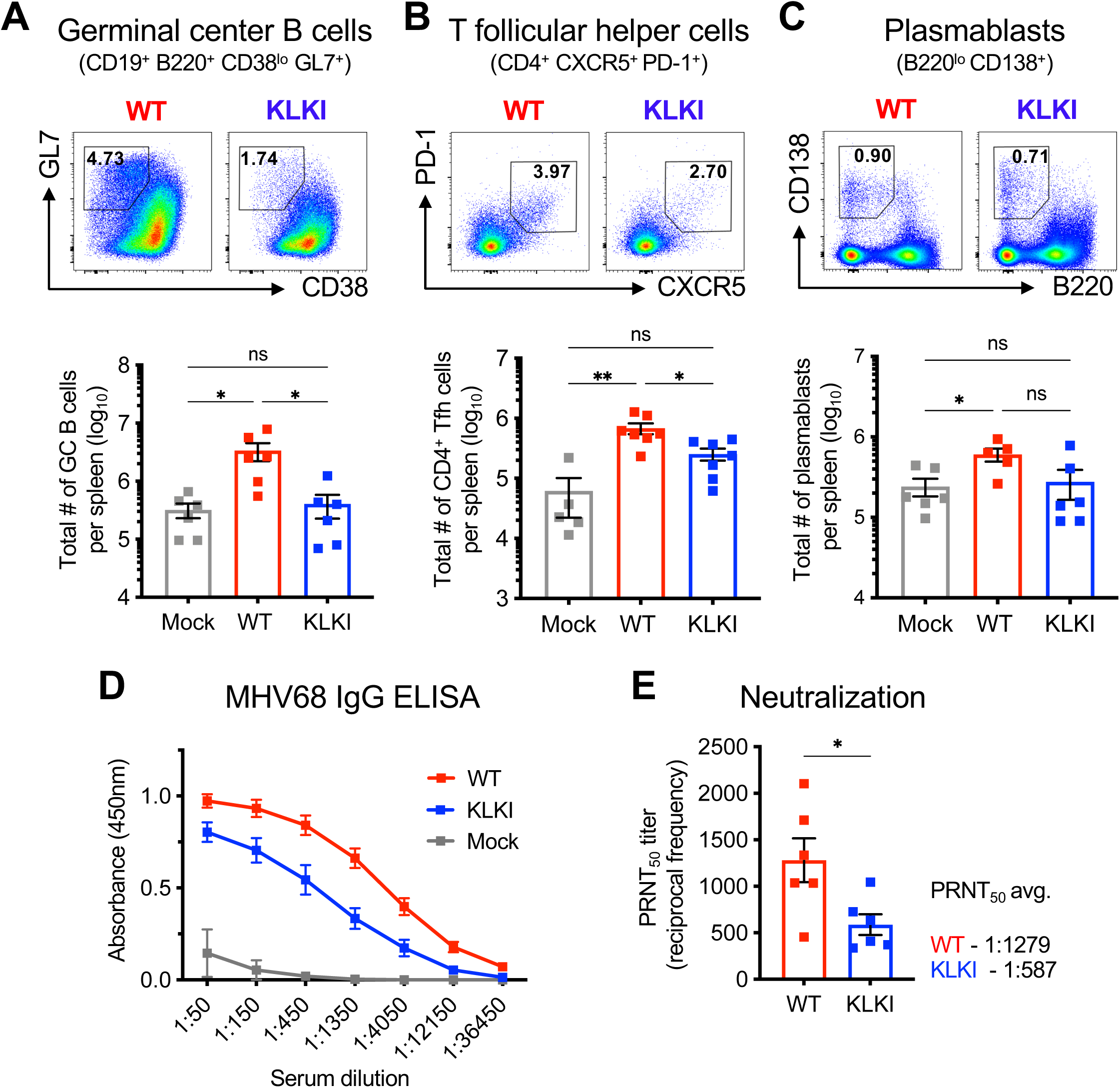
B cell activation and functional antibody responses are reduced following KLKI MHV68 infection. C57BL/6 mice were mock-infected or infected IN with 1000 PFU of the indicated virus. Mice were sacrificed on day 16-18 postinfection. Spleens from infected mice were analyzed via flow cytometry. Representative gates from each infection are shown (top) and total cell counts were quantified (bottom). (A) Germinal center B cells were defined as CD19^+^/B220^+^/CD38^lo^/GL7^+^, (B) T follicular helper cells as CD3e^+^/CD4^+^/CXCR5^+^/PD-1^+^, and (C) plasmablasts as B220^lo^/CD138^+^. (D and E) Serum was isolated from infected animals for downstream analysis. (D) MHV68-specific IgG was determined by ELISA. (E) Neutralizing capacity of MHV68-specific antibodies was determined by plaque-reduction neutralization test (PRNT) assays using the indicated concentrations of serum. PRNT_50_ titers were determined where each sample neutralizes 50% of plaques on an indicator monolayer relative to negative controls. Groups of 3-5 mice were used for each infection and analysis. Results are means of 2-3 independent infections. (A-C) Each dot represents one mouse. Error bars represent standard error of the mean. *, ** denotes p<0.05, p<0.01, respectively, in a one-way ANOVA with Tukey’s test. ns – not significant. (E) * denotes p<0.05 in an unpaired Student’s *t-*test.

To assess the antiviral impact of reduced B cell activation during KLKI MHV68 infection, we compared MHV68-specific humoral responses after WT and KLKI MHV68 infections. We quantified MHV68-specific IgG in mouse sera by ELISA, finding that virus-specific antibodies generated in response to KLKI MHV68 infection had a reduced capacity to recognize MHV68 lytic antigens **(Fig. 2D)**. To assess antibody functionality, plaque-reduction neutralization titer (PRNT) assays were conducted to determine the neutralizing capacity of virus-specific Ig. PRNT_50_ titers were 1:587 for KLKI-infected animals while WT MHV68 infection resulted in a PRNT_50_ titer of 1:1279, indicating that the humoral response to KLKI infection was ∼2.2-fold less potent than the response to WT MHV68 infection. **(Fig. 2E)**. Collectively, these data indicate that B cell activation and humoral immunity is less potent during latency establishment by KLKI MVH68 relative to WT virus.

Effector CD4+ and CD8+ T cell expansion in the spleen contributes to splenic inflammation; CD4+ T cells that produce IFNγ play a vital role in controlling MHV68 replication during chronic infection (24–26), and CD8+ T cells directly recognize and kill infected cells through perforin and granzyme production, while also secreting TNFα and IFNγ to drive antiviral inflammatory responses (27, 28). In flow cytometry-based comparisons of T cell activation following KLKI and WT MHV68 infection, effector CD4+ and CD8+ T cell compartments (CD4^+^/CD8^+^ CD44^hi^ CD62L^lo^) were much smaller in KLKI MHV68-infected animals **(Fig. 3A-B)**. However, viral antigen-specific effector CD8+ T cells were similarly induced by both viruses, as CD8+ T cells that recognized the p56 MHC class I tetramer containing the ORF6_487-495_ peptide were significantly elevated during infection with either virus **(Fig. 3C)**. Thus, despite reduced lytic replication and reactivation (**Fig. S2** and ref. (12)), KLKI MHV68 drives virus-specific T cell responses to an immunodominant MHV68 T cell epitope, which is consistent with data from replication defective viruses as well (18, 29). Overall, these data suggest that repression of lytic replication by kLANA during KLKI MHV68 infection correlates with diminished effector T cell activation, but not virus-specific effector CD8+ T cell expansion.

**Figure 3.**
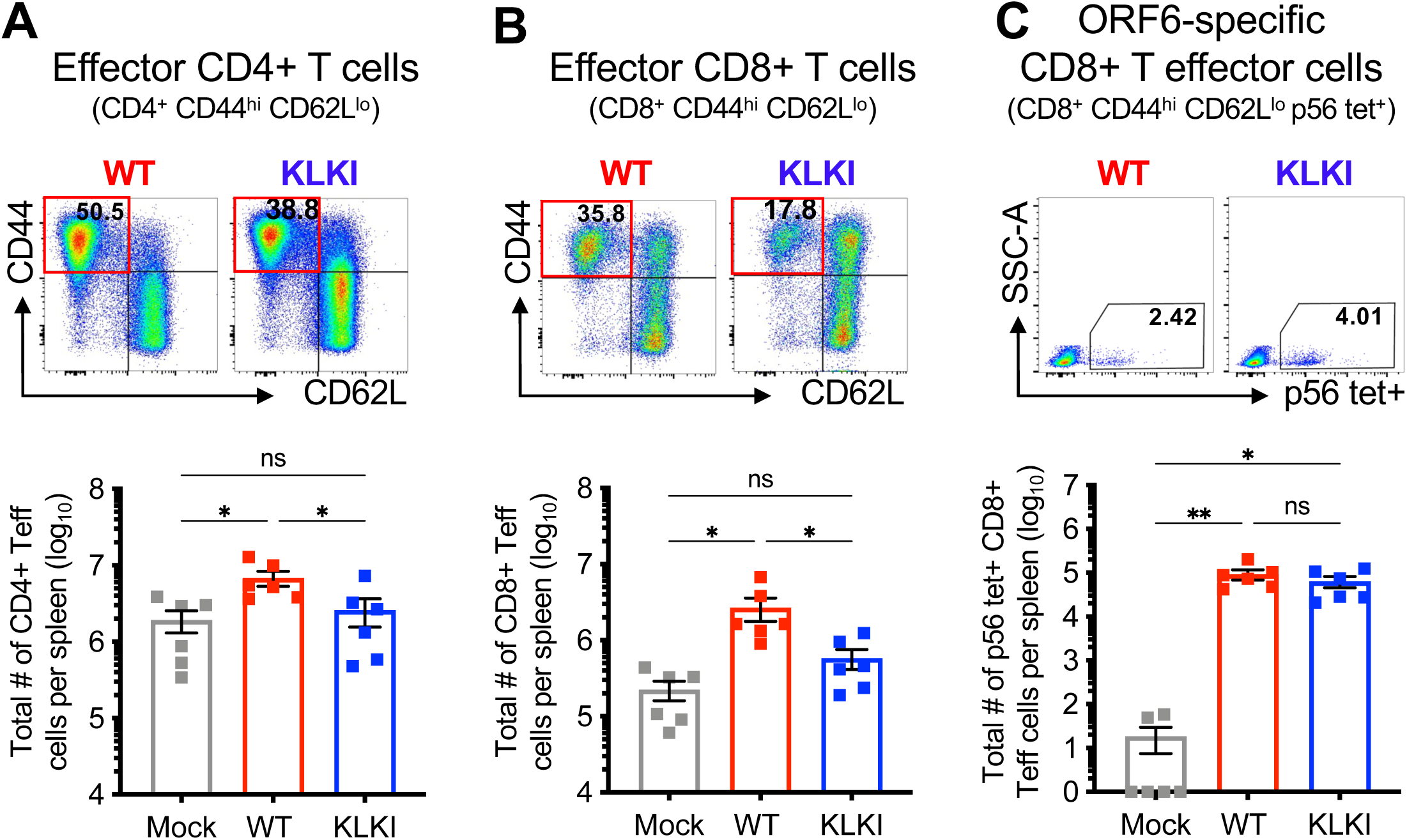
Despite limited T cell activation, virus-specific CD8+ T cell responses arise during KLKI MHV68 infection. C57BL/6 mice were mock-infected or infected IN with 1000 PFU of the indicated virus. Mice were sacrificed on day 16-18 postinfection. Spleens from infected mice were analyzed via flow cytometry. Representative gates from each infection are shown (top) and total cell counts were quantified (bottom). (A) Effector CD4+ T cells were gated as CD3e+/CD4+/CD44hi/CD62Llo, (B) effector CD8+ T cells as CD3e+/CD8+/CD44hi/CD62Llo, and (C) ORF6-specific effector CD8+ T cells as CD3e+/CD8+/CD44hi /CD62Llo/p56 tetramer+. Groups of 3-5 mice were used for each infection and analysis. Each dot represents one mouse. Results are means of 2-3 independent infections. Error bars represent standard error of the mean. *,** denotes p<0.05, p<0.01, respectively, in a one-way ANOVA with Tukey’s test. ns – not significant.

### Inoculating dose is not a major determinant of adaptive immune activation

While the infectious dose for KSHV in humans is not known, a dose-response analysis of MHV68 infection after intranasal infection found that inoculation with doses ranging from 0.1 to 400,000 PFU results in equivalent levels of latent infection in spleens of mice (30). However, the impact that the initial inoculating dose of any GHV has on immune activation is not defined. We reasoned that reduced lytic replication and reactivation by KLKI MHV68 could reduce the amount of danger signaling and antigen availability for potent immune activation, leading us to hypothesize that a high-dose inoculation of KLKI MHV68 infection would enhance immune activation and potency. We therefore performed a dose-response analysis for WT and KLKI MHV68 using infectious doses of 10, 1000, and 100,000 PFU and assessed adaptive immune responses.

Splenomegaly induced by KLKI MHV68 remained minimal, even following inoculation with 100,000 PFU, and enumeration of immune cell subsets in spleens indicated that the percentages of GC B cells, effector CD4+, and effector CD8+ T cells were significantly lower during KLKI MHV68 infection compared to WT virus infections for all doses tested **(Fig. 4A-D)**. While WT MHV68 infection stimulated a more potent IgG antibody response at both low and intermediate doses, virus-specific antibody responses were comparable between WT and KLKI MHV68 infection only for the 100,000 PFU dose **(Fig. 4E)**. Importantly, the frequencies of latently infected cells in spleens were comparable between both viruses at all doses tested **(Fig. 4F and Table 1)**. Note that evaluations for 10 PFU infections required analysis on day 28 post-infection, as virus was not present in spleens at the traditional day 16-18 timepoint (not shown). Together these data demonstrate that, although virus-specific antibody production increased in a dose-dependent manner for KLKI MHV68, general immune activation was predominantly independent of inoculating dose.

**Figure 4.**
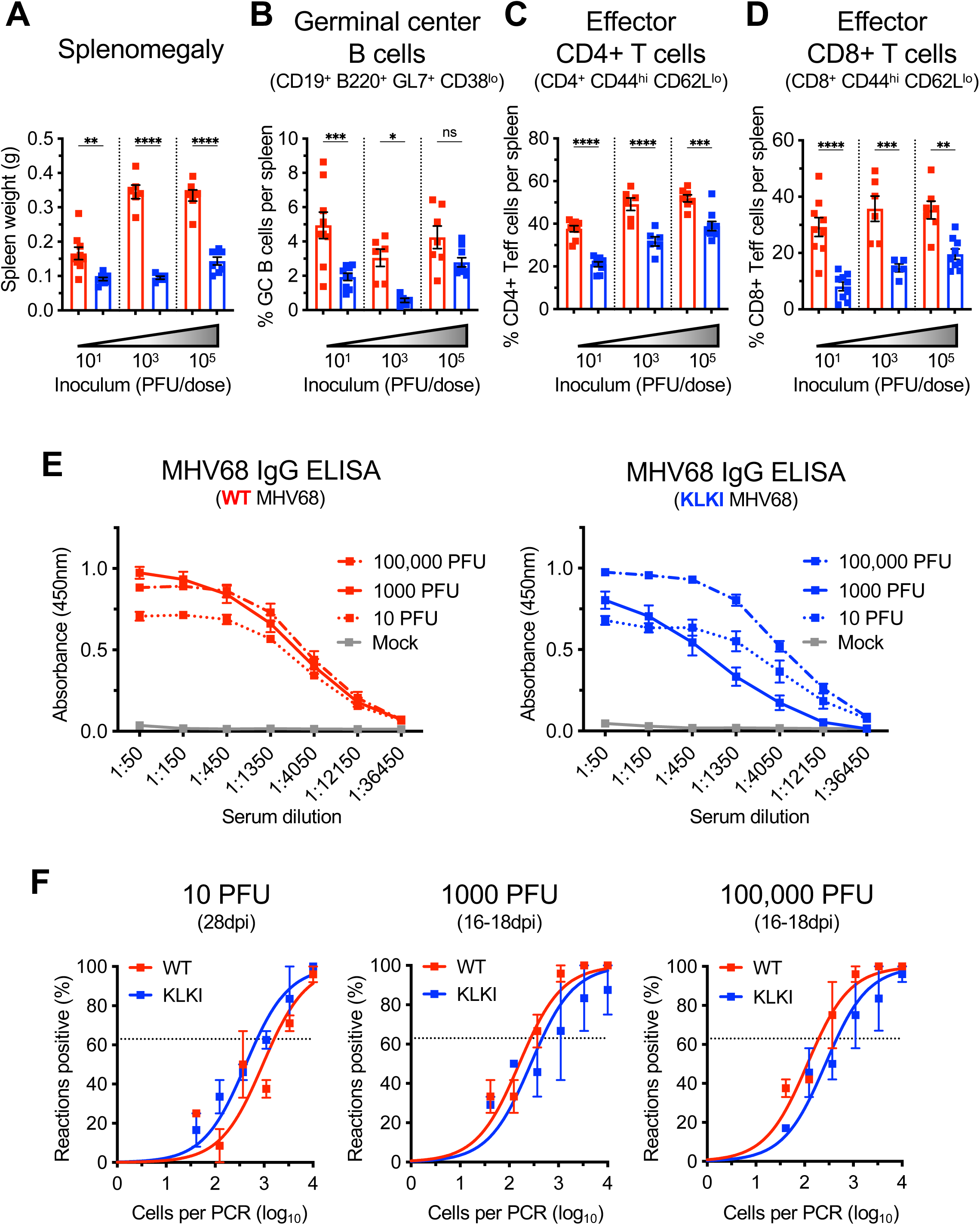
Adaptive immune activation in response to KLKI MHV68 infection is reduced regardless of infectious dose. C57BL/6 mice were infected IN with 10, 1000, or 100,0000 PFU of the indicated virus. Mice were sacrificed on day 16-18 postinfection (1000, or 100,000 PFU) or day 28 postinfection (10 PFU). (A) Spleens were weighed as an assessment of splenomegaly at each dose tested and the percentages of (B) germinal center B cells (CD19^+^/B220^+^/CD38^lo^/GL7^+^), (C) effector CD4+ T cells (CD3e^+^/CD4^+^/CD44^hi^/CD62L^lo^), and (D) effector CD8+ T cells (CD3e^+^/CD8^+^/CD44^hi^/CD62L^lo^) from infected spleens were analyzed and quantified via flow cytometry. (E) Serum was isolated from infected animals and analyzed in MHV68-specific IgG ELISAs for WT MHV68 (red; left) or KLKI MHV68 (blue; right) infections at each dose. (F) Single-cell suspensions of splenocytes were serially diluted, and the frequencies of cells harboring MHV68 genomes were determined using a limiting-dilution PCR analysis. Groups of 3-5 mice were used for each infection and analysis. Results are means of 2-3 independent infections. (A-D) Each dot represents one mouse. Error bars represent standard error of the mean. *,**,***,**** denotes p<0.05, p<0.01, p<0.001, p<0.0001, respectively, in a one-way ANOVA with Tukey’s test. ns – not significant.

**Table 1.**
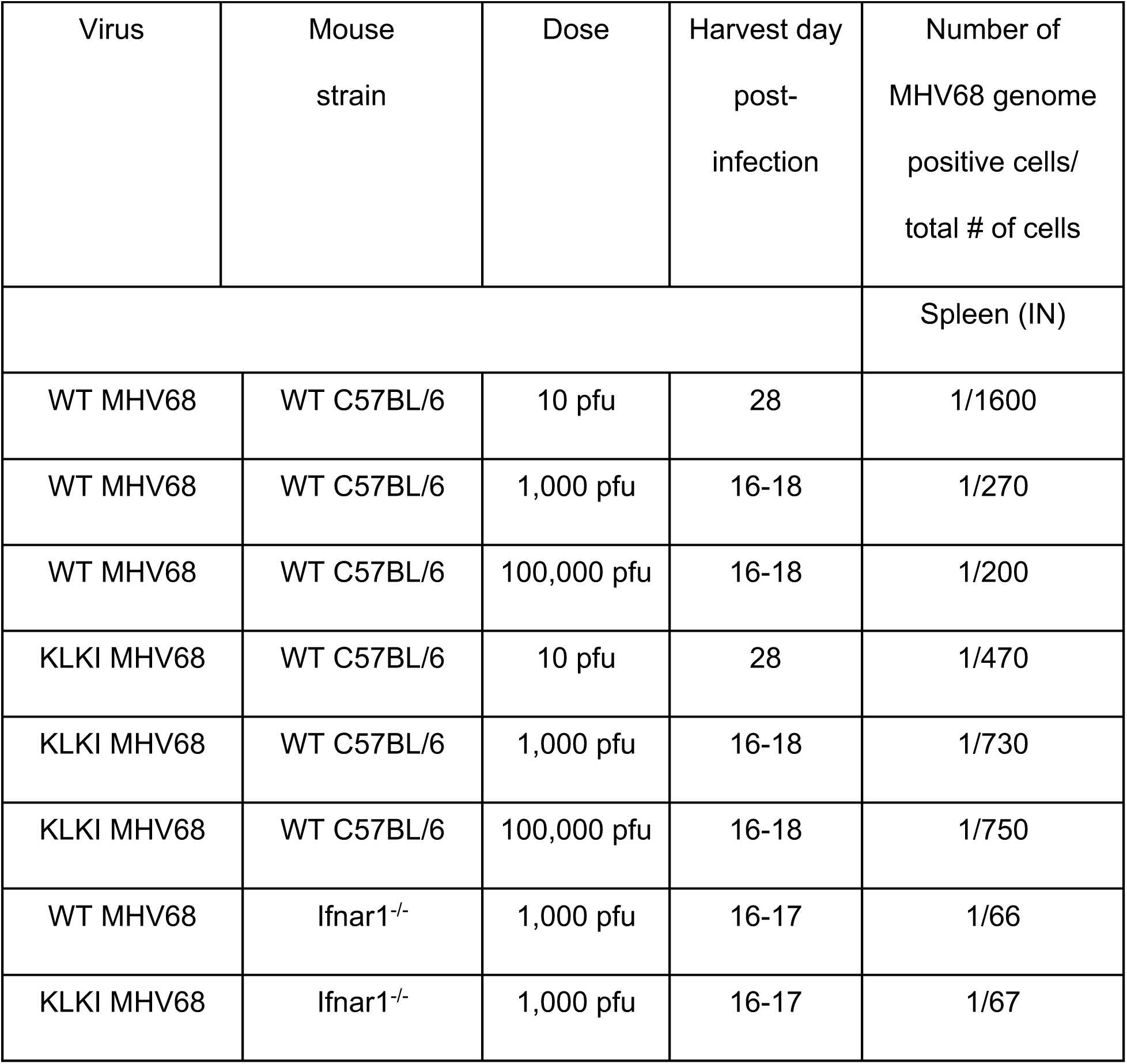
Frequencies of cells harboring MHV68 genomes in latently infected mice.

|  |  |  |  |  |
| --- | --- | --- | --- | --- |
| Virus | Mouse strain | Dose | Harvest day post-infection | Number of MHV68 genome positive cells/ total # of cells |
|  |  |  |  | Spleen (IN) |
| WT MHV68 | WT C57BL/6 | 10 pfu | 28 | 1/1600 |
| WT MHV68 | WT C57BL/6 | 1,000 pfu | 16-18 | 1/270 |
| WT MHV68 | WT C57BL/6 | 100,000 pfu | 16-18 | 1/200 |
| KLKI MHV68 | WT C57BL/6 | 10 pfu | 28 | 1/470 |
| KLKI MHV68 | WT C57BL/6 | 1,000 pfu | 16-18 | 1/730 |
| KLKI MHV68 | WT C57BL/6 | 100,000 pfu | 16-18 | 1/750 |
| WT MHV68 | lfnar1<sup>-/-</sup> | 1,000 pfu | 16-17 | 1/66 |
| KLKI MHV68 | lfnar1<sup>-/-</sup> | 1,000 pfu | 16-17 | 1/67 |

### Abrogation of type I IFN signaling enhances productive viral replication during KLKI MHV68 infection

Since high-dose KLKI MHV68 infection did not promote activation of the adaptive immune response to the same extent as WT MHV68 infection, we reasoned that a lytic replication threshold, rather than a generic antigenic load, needs to be breached to potently activate adaptive immune responses during MHV68 infection. We therefore sought to enhance KLKI lytic replication to evaluate the impact on adaptive immune stimulation. MHV68 lytic replication is controlled by type I interferon (IFN) responses (31, 32). Thus, we hypothesized that KLKI MHV68 lytic replication would be enhanced in type I IFN non-responsive cells in a manner that surpasses the immune engagement threshold.

As an initial test, we utilized CRISPR-Cas9 gene editing to ablate the alpha subunit of the type I IFN receptor (*Ifnar1*) in 3T12 fibroblasts, rendering them type I IFN nonresponsive, and evaluated KLKI MHV68 infection relative to WT virus **(Fig. 5)**. *Ifnar1* ablation was confirmed by flow cytometry **(Fig. 5A)** and lack of STAT1 phosphorylation (pSTAT1) after treatment with recombinant IFN-β **(Fig. 5B)**. We then infected *Ifnar1*-deficient fibroblasts with either WT or KLKI MHV68 and evaluated viral transcription over time. While control fibroblasts infected with KLKI MHV68 exhibited low-level lytic transcription that did not appear to amplify to the same extent as WT virus infections, both viruses readily progressed through the lytic cascade following infection of *Ifnar1*-deficient cells **(Fig. 5D).** Immunoblot and immunofluorescence analyses revealed that lytic viral gene products and plaque size were dramatically increased, respectively, following KLKI MHV68 infection of *Ifnar1*-deficient cells relative to mSAFE non-target control cells **(Fig. 5E-F)**. Consistent with the observation that lytic viral gene transcription and translation were enhanced in the absence of type I IFN signaling, WT and KLKI MHV68 replicated more efficiently in *Ifnar1*-deficient fibroblasts **(Fig. 5C)**. While still reduced compared to WT MHV68, a phenotype we attribute to continued kLANA-mediated suppression of *ORF50*/RTA despite IFN non-responsiveness, these results demonstrate that *Ifnar1* deletion potently enhances viral antigen accumulation and viral replication during KLKI MHV68 infection.

**Figure 5.**
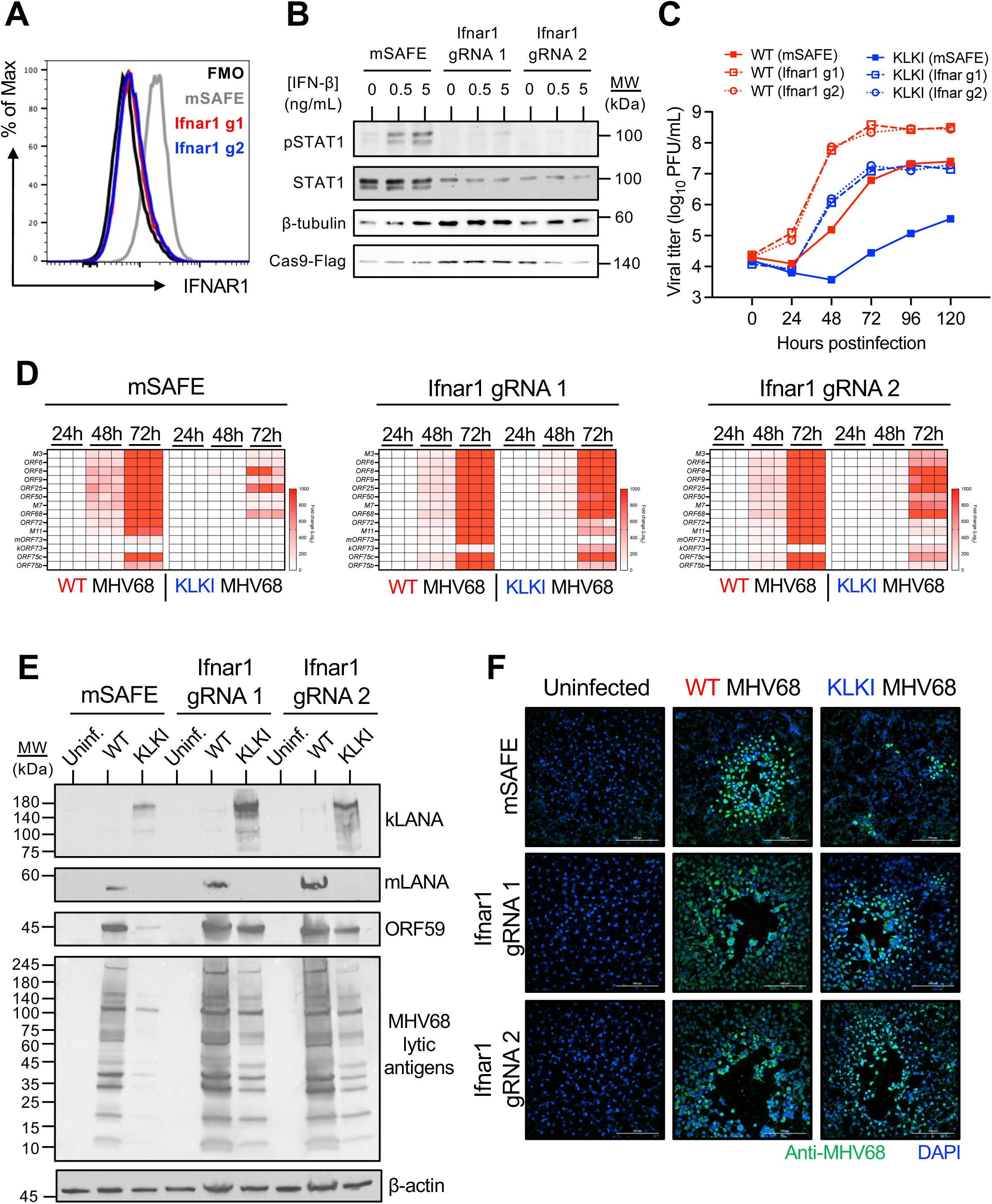
*Ifnar1* ablation in murine fibroblasts promotes lytic antigen accumulation and viral replication during KLKI MHV68 infection. (A) *Ifnar1*-deficient or non-target control (mSAFE) 3T12 fibroblast pools were stained with *Ifnar1*-specific antibodies and analyzed by flow cytometry. (B) Cells were treated with recombinant IFN-β at indicated doses to determine efficacy of *Ifnar1*-specific deletion. Lysates were collected 2.5 hours after treatment and immunoblot analyses were performed using antibodies to detect indicated proteins. (C-D,F) Cells were infected with the indicated virus at an MOI of 0.1 PFU/cell. (C) Viral titers were determined at the indicated times postinfection by plaque assay. (D) RNA was collected at the indicated times postinfection. Viral transcripts were quantified by qRT-PCR relative to *B2m* using the ΔΔCT method and normalized to transcript abundance at 24 hours postinfection. (E) Cells were infected at an MOI of 1 PFU/cell and lysates were collected 48 hours postinfection. Immunoblot analyses were performed using antibodies that recognize the indicated proteins. (F) Cells were fixed 72 hours postinfection and stained for indirect immunofluorescence analysis using antibodies that recognize the indicated proteins. DNA was stained with DAPI. Scale bar – 200um.

To test the immune engagement threshold hypothesis *in vivo*, we infected *Ifnar1^-/-^*C57BL/6 mice with WT and KLKI MHV68 and evaluated viral replication, latency, and adaptive immunity. Consistent with previous studies (33), after IN inoculation WT MHV68 viral titers in lungs were approximately 100-fold higher in *Ifnar1*^-/-^ mice than WT C57BL/6 mice at day 7 post-infection and exhibited delayed clearance on day 10 post-infection **(Fig. 6A)**. This was also the case for KLKI MHV68. Although still reduced compared to WT virus, splenomegaly did occur following KLKI MHV68 infection of *Ifnar1*^-/-^ mice with spleen weights significantly higher than uninfected controls **(Fig. 6B)**. As in WT C57BL/6 mice (see **Fig. 4E**), both KLKI and WT MHV68 established latency equivalently in spleens of *Ifnar1*^-/-^ mice on day 16-17 post-infection **(Fig. 6C and Table 1).** While KLKI MHV68 reactivation from latency remained reduced compared to WT virus, both WT and KLKI MHV68 exhibited an increased frequency of reactivating cells in comparison to WT C57BL/6 mice and no persistent viral replication was observed **(Fig. 6D**, **Table 2 and Fig. S2)**. These data indicate that the absence of type I IFN signaling in *Ifnar1^-/-^* mice enhances KLKI MHV68 lytic replication and reactivation and restores splenomegaly.

**Figure 6.**
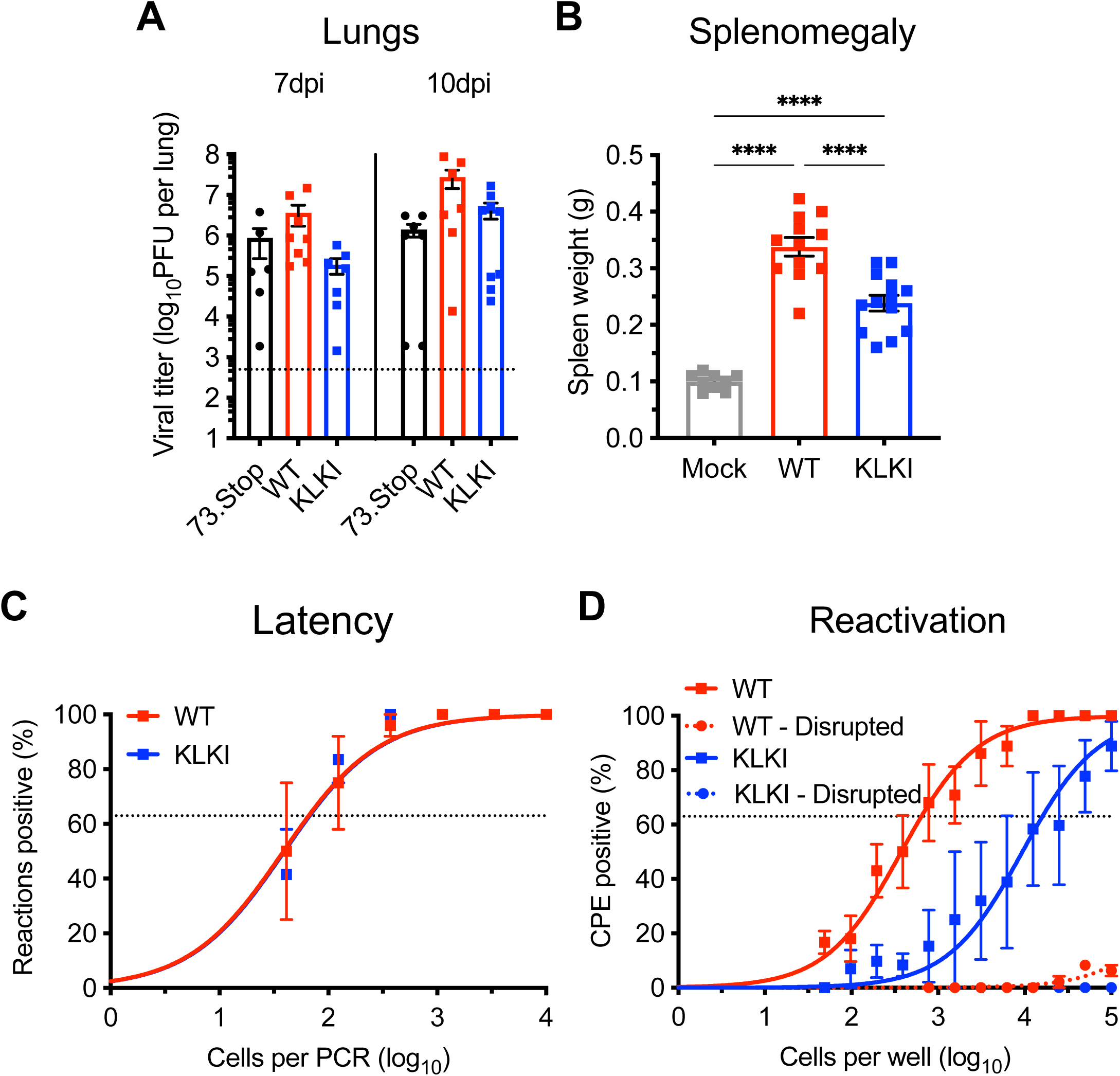
Lytic replication is enhanced during KLKI MHV68 infection of *Ifnar1^-/-^* mice and promotes splenomegaly. *Ifnar1^-/-^* mice were mock-infected or infected IN with 1000 PFU of the indicated virus. (A) Mice were sacrificed on day 7 or 10 postinfection. Lungs were homogenized and viral titers were determined by plaque assay. (B-D) Mice were sacrificed on day 16-17 postinfection. (B) Spleens were weighed as an assessment of splenomegaly. (C) Single-cell suspensions of splenocytes were serially diluted, and the frequencies of cells harboring MHV68 genomes were determined using a limiting-dilution PCR analysis. (D) Reactivation frequencies were determined by *ex vivo* plating of serially diluted splenocytes on an indicator monolayer. Cytopathic effect was scored one week postplating. Groups of 3-5 mice were used each infection and pooled for analysis (C-D). Results are means of 2-3 independent infections. Error bars represent standard error of the mean. ***,**** denotes p<0.001, p<0.0001, respectively, in a one-way ANOVA with Tukey’s test. ns – not significant.

**Table 2.**
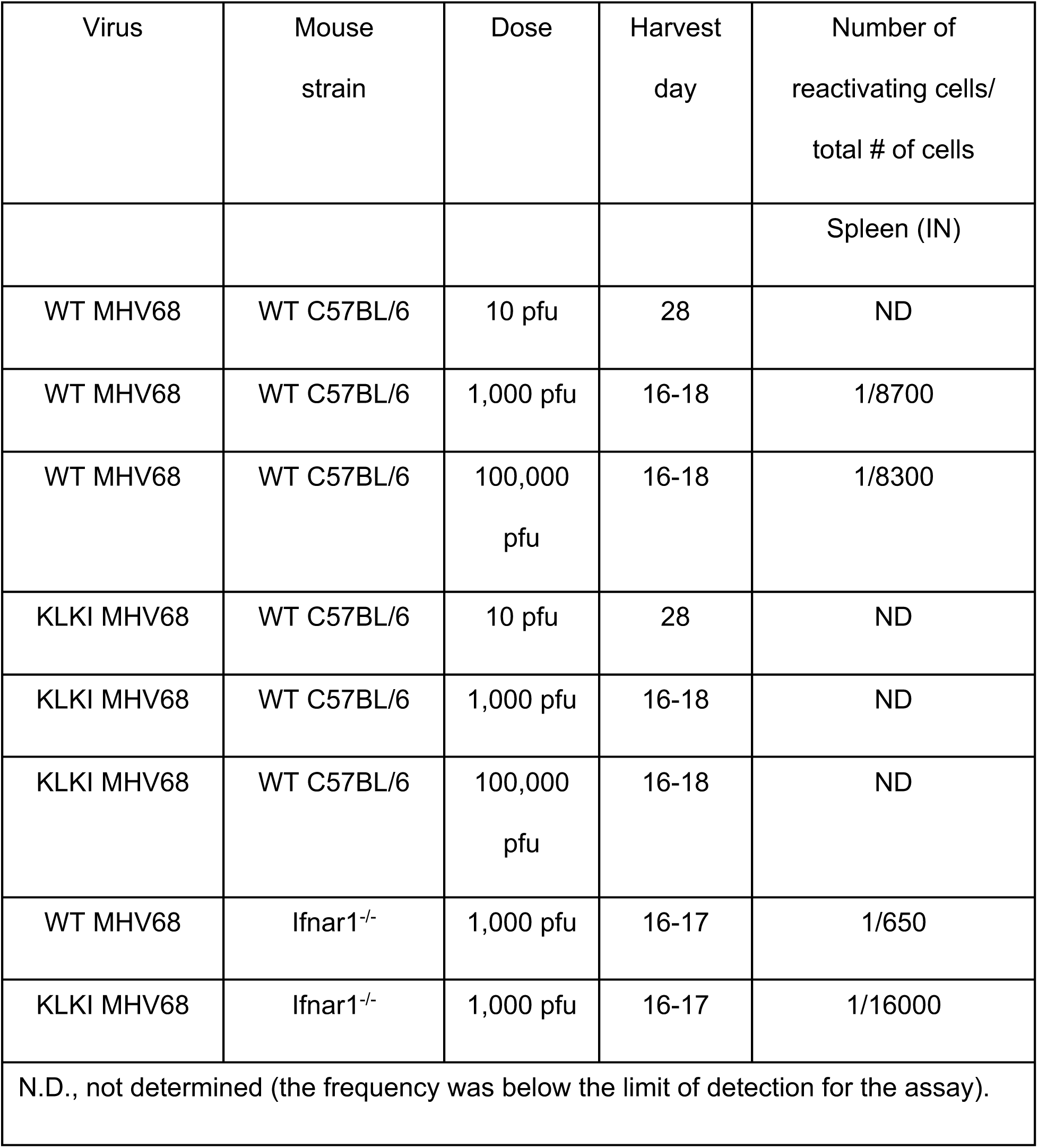
Frequencies of latently infected cells reactivating ex vivo.

| Virus | Mouse strain | Dose | Harvest day | Number of reactivating cells/<br>total # of cells |
| --- | --- | --- | --- | --- |
|  |  |  |  | Spleen (IN) |
| WT MHV68 | WT C57BL/6 | 10 pfu | 28 | ND |
| WT MHV68 | WT C57BL/6 | 1,000 pfu | 16-18 | 1/8700 |
| WT MHV68 | WT C57BL/6 | 100,000<br>pfu | 16-18 | 1/8300 |
| KLKI MHV68 | WT C57BL/6 | 10 pfu | 28 | ND |
| KLKI MHV68 | WT C57BL/6 | 1,000 pfu | 16-18 | ND |
| KLKI MHV68 | WT C57BL/6 | 100,000<br>pfu | 16-18 | ND |
| WT MHV68 | lfnar1<sup>-/-</sup> | 1,000 pfu | 16-17 | 1/650 |
| KLKI MHV68 | lfnar1<sup>-/-</sup> | 1,000 pfu | 16-17 | 1/16000 |
| N.D., not determined (the frequency was below the limit of detection for the assay). |  |  |  |  |

### KLKI MHV68 infection stimulates potent antiviral adaptive immunity in *Ifnar1^-/-^* mice

Since MHV68-induced splenomegaly is dependent on T cell activation (14, 34), enhanced splenomegaly following infection of *Ifnar1*^-/-^ mice with KLKI MHV68 suggested that adaptive immune activation occurred in this model. Indeed, in contrast to findings in WT C57BL/6 mice, KLKI MHV68 infection caused potent GC B cell expansion in *Ifnar1*^-/-^ mice **(Fig. 7A)** that correlated with increased Tfh cell expansion when compared to mock-infected spleens **(Fig. 7B)**. Additionally, both WT and KLKI MHV68 infections caused a 10-fold increase in plasmablast expansion relative to uninfected controls **(Fig. 7C)**. In agreement with the B cell activation phenotypes, serum collected from both WT and KLKI MHV68 infected *Ifnar1^-/-^* mice contained high levels of MHV68-specific IgG, and both sera exhibited comparable capacities to neutralize MHV68 in PRNT_50_ assays **(Fig. 7D-E)**.

**Figure 7.**
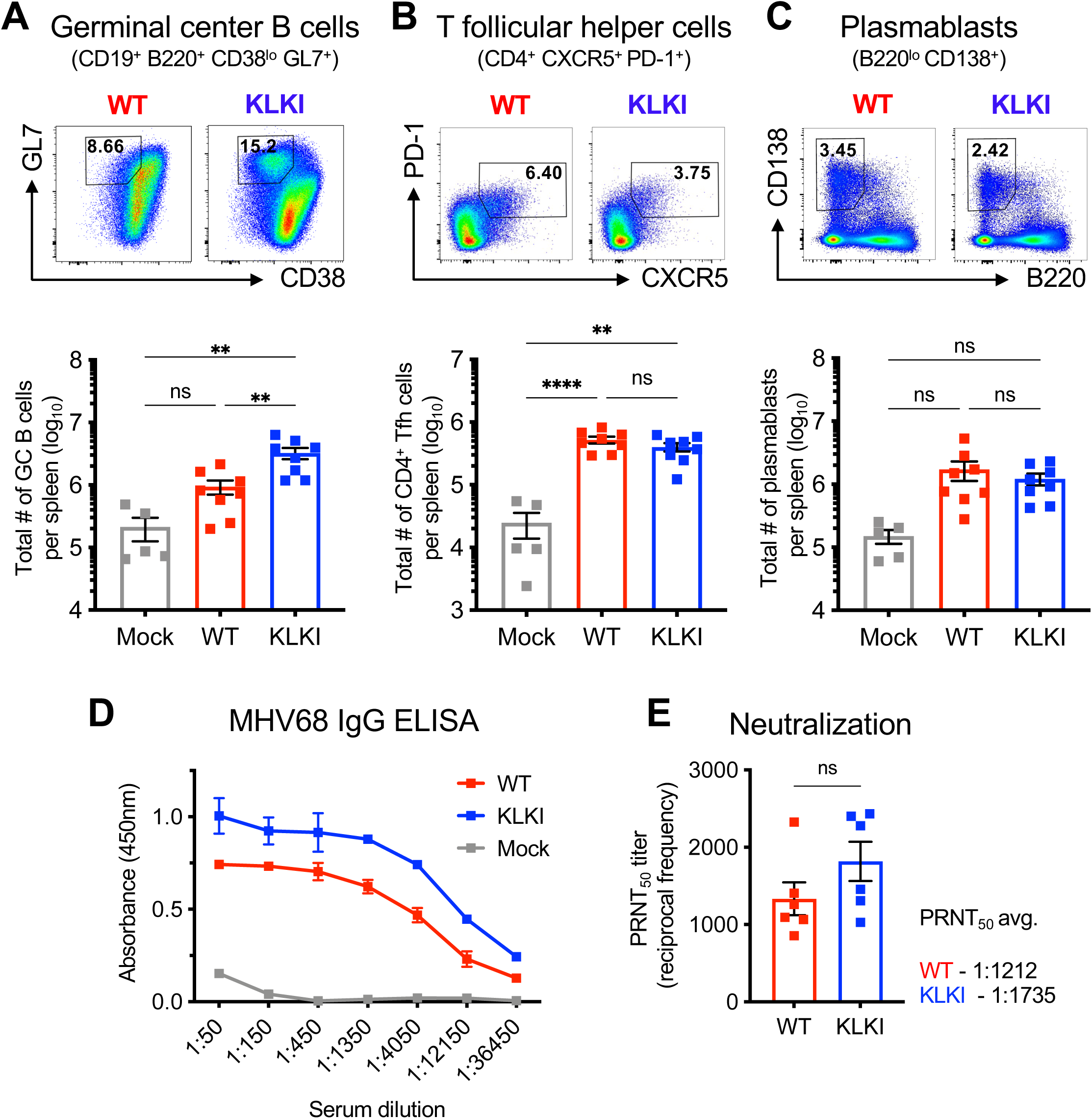
B cell activation and antibody responses are enhanced during KLKI MHV68 infection of *Ifnar1^-/-^* mice. *Ifnar1^-/-^* mice were infected IN with 1000 PFU of the indicated virus. Mice were sacrificed on day 16-17 postinfection. Spleens from infected mice were analyzed via flow cytometry. Representative gates from each infection are shown (top) and total cell counts were quantified (bottom). (A) Germinal center B cells were defined as CD19^+^/B220^+^/CD38^lo^/GL7^+^, (B) T follicular helper cells as CD3e^+^/CD4^+^/CXCR5^+^/PD-1^+^, and (C) plasmablasts as B220^lo^/CD138^+^. (D and E) Serum was isolated from infected animals for downstream analysis. (D) MHV68-specific IgG was quantified by ELISA. (E) Neutralizing capacity of MHV68-specific antibodies was determined by plaque-reduction neutralization test (PRNT) assays using the indicated concentrations of serum. PRNT_50_ titers were determined where each sample neutralizes 50% of plaques on an indicator monolayer relative to negative controls. Groups of 3-5 mice were used for each infection and analysis. Results are means of 2-3 independent infections. (A-C) Each dot represents one mouse. Error bars represent standard error of the mean. **,**** denotes p<0.01, p<0.0001, respectively, in a one-way ANOVA with Tukey’s test. ns – not significant.

Consistent with splenomegaly occurring following KLKI MHV68 infection of *Ifnar1^-/-^* mice, effector CD4+ T cell activation in spleens was similar for both WT and KLKI MHV68 **(Fig. 8A)**. Effector CD8+ T cell populations also were similarly elevated following infection with either virus **(Fig. 8B)**, with equivalent numbers of p56 tetramer-specific effector CD8+ T cells elicited **(Fig. 8C)**. Thus, KLKI MHV68 infection of *Ifnar1^-/-^* mice promotes B and T cell responses that are comparable to WT MHV68 infection. Taken together, these results suggest that the level of lytic replication/reactivation during MHV68 infection is a primary driver of adaptive immune engagement. Moreover, these findings provide evidence that suppression of lytic viral transcription and replication by kLANA functions in immune evasion, enabling efficient latency establishment without potent adaptive immune stimulation.

**Figure 8.**
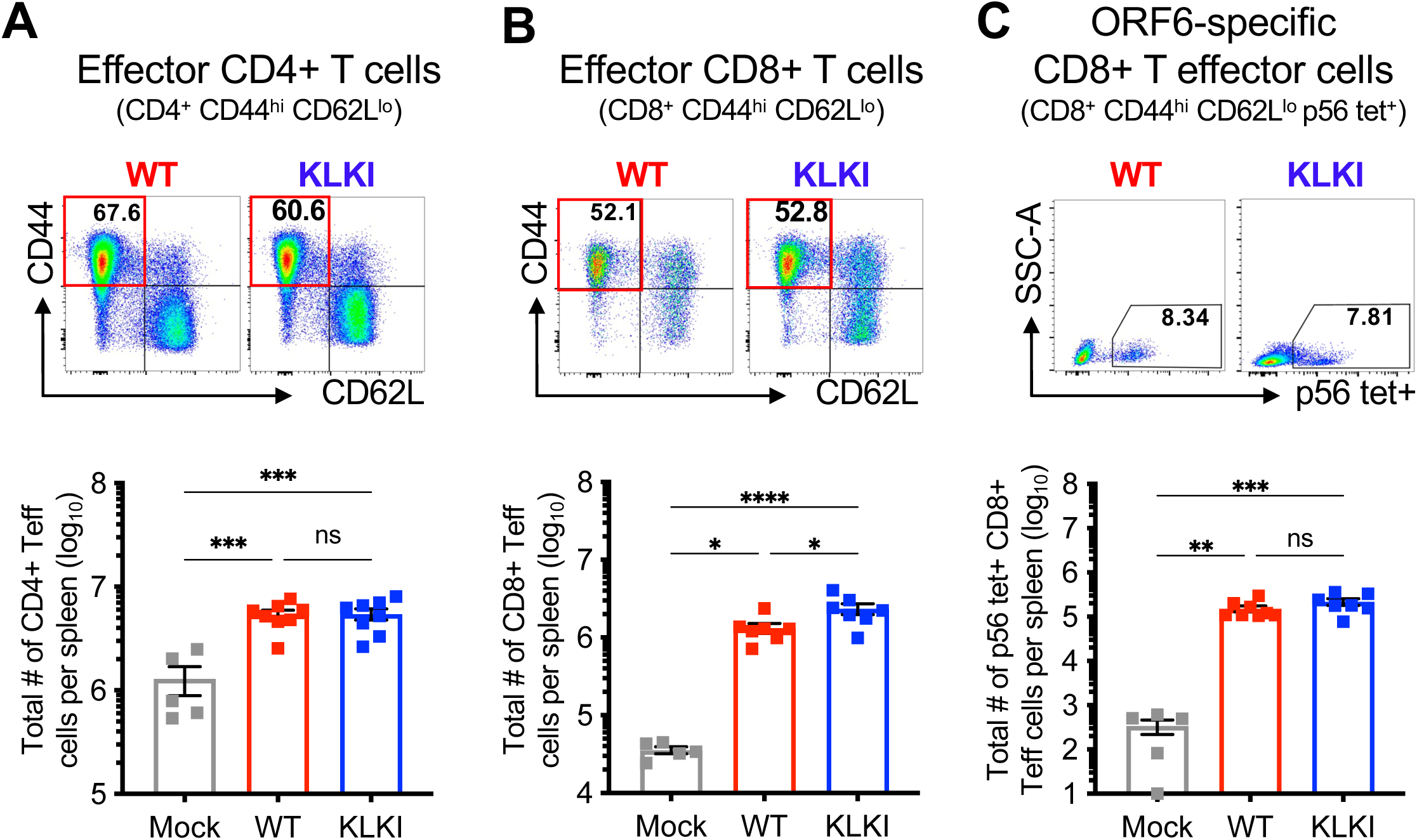
T cell activation and virus-specific effector CD8+ T cell responses are enhanced following KLKI MHV68 infection of *Ifnar1^-/-^* mice. *Ifnar1^-/-^* mice were mock-infected or infected IN with 1000 PFU of the indicated virus. Mice were sacrificed on day 16-17 postinfection. Spleens from infected mice were analyzed via flow cytometry. Representative gates from each infection are shown (top) and total cell counts were quantified (bottom). (A) Effector CD4+ T cells were gated as CD3e^+^/CD4^+^/CD44^hi^/CD62L^lo^, (B) effector CD8+ T cells as CD3e^+^/CD8^+^/CD44^hi^/CD62L^lo^, and (C) ORF6-specific effector CD8+ T cells as CD3e^+^/CD8^+^/CD44^hi^ /CD62L^lo^/p56 tetramer^+^. Groups of three to five mice were used for each infection and analysis. Each dot represents one mouse. Results are means of 2-3 independent infections. Error bars represent standard error of the mean. *,**,***,**** denotes p<0.05, p<0.01, p<0.001, p<0.0001, respectively, in a one-way ANOVA with Tukey’s test. ns – not significant.

## DISCUSSION

Lytic replication facilitates GHV dissemination and transmission to new hosts; however, this process is highly immunogenic, promoting the detection and neutralization of infected cells by host immune responses. By switching to latency, GHVs minimize viral gene expression to factors that facilitate viral episome persistence and host cell survival, thereby evading host immunity. Here we demonstrate that an MHV68 virus expressing kLANA in place of mLANA exhibits much reduced viral gene expression relative to WT and mLANA-deficient MHV68, suggestive of a pro-latent phenotype. Despite establishing latency at frequencies equivalent to WT MHV68, virus-specific adaptive immune responses were comparatively much lower following KLKI MHV68 infection of mice. Indeed, reductions in B and T cell activation during KLKI virus infections correlated with a reduction in splenomegaly, suggesting that proinflammatory responses during latency establishment are reduced when lytic replication and/or reactivation is suppressed. IFN responses act as host restriction factors to viral replication, and deletion of the *Ifnar1* receptor enhances lytic MHV68 replication/reactivation. Consistent with this, infection of *Ifnar1^-/-^*mice resulted in enhanced KLKI MHV68 lytic replication and reactivation, infection-induced splenomegaly, and adaptive immune cell activation *in vivo*. Since enhanced lytic replication/reactivation during KLKI MHV68 infections coincided with stronger immune cell activation in *Ifnar1^-/-^* mice, we propose that a lytic replication threshold must be surpassed for potent adaptive immune cell activation in response to GHV infection.

It is important to note that we currently include reactivation within lymphoid organs in our overarching interpretation as it is challenging to separate the immune activating impacts of reactivation *in situ* from primary acute replication. The rationale for this interpretation is primarily based on the observation that KLKI MHV68 exhibits increased reactivation efficiency following infection of *Ifnar1^-/-^* mice. We attempted to address the contribution of reactivation more directly through conditional deletion of mRTA-encoding *ORF50 in vivo* as previously described (35). Although we previously reported reduced splenomegaly and reactivation relative to WT virus (35), impacts on immune activation following infection of CD19-Cre mice with floxed *ORF50* MHV68 were variable and inconclusive and therefore did not provide clarity (not shown). The converse argument that reactivation does not influence immune activation could also be made given that following infection with 10 PFU of either WT or KLKI MHV68, reactivation remained below the limit of detection for *both* viruses, yet differences in adaptive immune activation were still observed. This seems to suggest that additional factors beyond measurable reactivation may drive these phenotypes. Suffice it to say, whether reactivation contributes directly to the phenotypes observed here remains unresolved.

LANA homologs mediate viral episome maintenance and segregation within latently infected cells. In addition to its essential functions in maintaining latent viral genomes, kLANA also represses *ORF50*/RTA transcription and subsequent lytic replication soon after naked viral DNA is deposited in host cells upon *de novo* infection (36). Indeed, kLANA recruits repressive chromatin modifying enzymes to the KSHV genome further suppressing viral transcription, as well as regulating host cell transcription (11, 37–40). Coupled with the facts that lytic gene expression is transient and tightly regulated during *de novo* KSHV infection (11, 36) and that KSHV primarily remains latent in a variety of cell types in culture (41), we hypothesize that this kLANA function evolved as a mechanism to limit immune detection by rapidly enforcing latency.

We previously demonstrated that kLANA, but not mLANA, could repress the promoter for the MHV68 *ORF50*/RTA gene in a manner that correlated with reduced *Orf50* transcription during infection (12), an observation that highlights a functional difference in these two LANA homologs. It is important to note that the transcription profile of KLKI MHV68 appears to reflect a repressive phenotype, and not just the absence of an mLANA-specific function, since mLANA-null MHV68 exhibited progressive amplification of viral gene expression over time, whereas KLKI MHV68 did not. It also is notable that repressed transcription during KLKI virus infections was overcome by expression of mRTA *in trans*, which further supports the interpretation that kLANA-mediates repression of lytic MHV68 transcription. Altogether, these data support the conclusion that kLANA-directed curtailment of viral transcription and replication skews KLKI MHV68 toward a more pro-latency phenotype, rather than undergoing the vigorous lytic replication exhibited by WT virus.

By extending our observations to KSHV and humans, we speculate that repression of lytic replication by kLANA enables primary KSHV infections to be minimally immune stimulating, hypothetically limiting both viral antigens and infection-associated inflammation. Indeed, after *de novo* infection of cells in culture, KSHV lytic gene transcription is transient, typically shutting off within ∼24 hours, and infectious virus particles are usually not detected (11, 42). Taken together with transcriptional analyses of KLKI MHV68 infection *in vitro*, the observation that adaptive immune cell activation and splenomegaly is reduced relative to mice infected with WT virus indicates that KLKI MHV68 infection is less immunostimulatory *in vivo*. Moreover, the absence of the inflammation-driven infectious mononucleosis (IM)-like disease splenomegaly implicates kLANA in dampening infection-associated pathologies.

To elaborate further on this idea, primary infection of humans by the related GHV, EBV, can be symptomatic, manifesting as IM, a pathology driven by inflammatory CD4+ and CD8+ T cells (43). The IM-like syndrome caused by MHV68 in mice also correlates with potent Th1-type virus-specific adaptive immunity after primary infection (44). Unlike EBV/MHV68, there are no known clinical symptoms associated with primary KSHV infection, suggesting that KSHV employs a “below the radar” approach to colonizing its hosts despite utilizing similar routes of transmission (45, 46). While other viral and host mechanisms also likely contribute, this highlights a putative role for kLANA in minimizing processes that drive immune stimulation during primary infection *in vivo*. These observations may also highlight a dichotomy between primary infections by MHV68 and EBV versus those of KSHV, arguing that MHV68 and EBV evolved to be inflammatory and immune activating to facilitate colonizing the host, whereas KSHV prefers a stealthier invasion tactic.

Since suppression of the lytic cycle correlated with reduced adaptive immune activation by KLKI MHV68, we next sought to determine if increasing the amount of viral antigens would overcome kLANA-mediated impacts on immune activation, starting by simply testing the dose-response of inoculating virus. The minimum infectious dose for MHV68 to establish latency in a mouse is less than a single PFU of virus (30), however the impact of inoculating dose on adaptive immunity to MHV68 has not to our knowledge been tested previously. In fact, how inoculating dose influences adaptive immunity against most viral infections in general is only minimally explored. We found that both B cell and T cell activation remained low for all doses of KLKI MHV68 tested relative to WT virus. Only virus-specific antibody production appeared to correlate with inoculating dose, which seems to agree with vaccination studies using non-replicating virus (18). Given that inoculating dose had no effect on KLKI MHV68 latency establishment, our data suggest that a kLANA-intrinsic function, and not simply antigenic input, reduces immune cell activation during infection.

Several possibilities could underlie the observation that high dose KLKI MHV68 infection is less immunogenic: 1) Despite a higher inoculating dose of antigen, the decreased production of lytic viral antigens within cells results in less antigen presentation for the host to detect, 2) fewer danger-associated molecular patterns are generated by KLKI MHV68 infection to elicit potent adaptive activation, or 3) kLANA modulates host-cell signaling pathways to prevent detection and/or activation by immune cells. These hypotheses, which are not mutually exclusive, could be tested in future experiments.

Since high-dose infections did not elicit strong immune activation, we next tested a model with enhanced viral replication to attempt to overcome kLANA-mediated dampening of adaptive immunity. IFNs act as critical mediators of host restriction during GHV infection, limiting acute replication and reactivation (31). Further, IFNα/β levels remain elevated during chronic GHV infection, implying that type I IFNs also regulate viral latency (47). Since enhancement of KLKI MHV68 lytic viral replication/reactivation in *Ifnar1^-/-^* mice led to increased adaptive immune stimulation and splenomegaly, we postulate that the absence of type I IFN signaling enables KLKI infections to surpass an antigenic or danger signaling threshold necessary to engage the adaptive immune response. These data suggest the existence of a trigger point, and once GHV lytic replication amplifies beyond this point during infection, potent antiviral adaptive immune responses are elicited. Although we cannot discount the idea that kLANA may work in concert with IFNα/β-signaling to suppress viral transcription and lytic replication, it is clear that enhancement of lytic viral replication/reactivation during infections of *Ifnar1^-/-^* mice is sufficient to potently stimulate adaptive immune cell activation regardless of whether kLANA-mediates repression of mRTA in this setting.

In essence, our data suggest that the “stealth” approach to infection adopted by KSHV, in part mediated by kLANA, likely enables viral colonization of the host while remaining below the triggering threshold that sounds the alarm for immune cell activation during primary infection. In agreement with this idea, it is interesting to note that KSHV seroconversion occurs several months after viral genomes are detectable in peripheral blood of newly infected individuals (48). By coupling low-level lytic replication at minimal infectious doses, a virus that is “pro-latent,” like KLKI MHV68 or KSHV, may evade eliciting potent antiviral adaptive immune responses after primary infection, while still efficiently colonizing a host.

Our work highlights an important aspect of studying KSHV-MHV68 chimeras to assess how viral factors affect GHV pathogenesis and development of host antiviral immunity. In general, information regarding how KSHV adaptive immune responses develop after primary infection in humans is lacking. Utilizing GHV chimeras like KLKI MHV68 provide an opportunity to elucidate mechanisms underlying KSHV pathogenesis in a living organism. Since our work demonstrates that KLKI MHV68 efficiently establishes latent infection while minimally stimulating adaptive immunity, we propose that kLANA-mediated repression of lytic viral replication similarly contributes to KSHV immune evasion in humans. Since humans can be infected simultaneously with multiple strains of KSHV (49, 50) and KSHV shows evidence of intra-strain recombination (51, 52), the absence of potent antiviral immunity could enable these events to occur, facilitating KSHV adaptation and evolution.

## MATERIALS AND METHODS

### Mice and infections

Male and female, C57BL/6 or *Ifnar1^-/-^*(B6(Cg)-Ifnar1tm1.2Ees/J) mice were purchased from Jackson Laboratories or were bred and maintained in house in sterile conditions. 6-to 12-week-old mice were anesthetized using isoflurane and inoculated with indicated dose of virus diluted in antibiotic-free DMEM (20 μL) for intranasal inoculations. Lungs, spleens, and serum were harvested at indicated times postinfection, as described previously (53).

### Cells and viruses

Vero-cre cells (54), 3T12 fibroblasts (ATCC CCL-164), Baby hamster kidney fibroblasts (BHK21) (ATCC CCL-10), and HEK 293T cells (ATCC CRL-3216) cells were cultured in Dulbecco’s Modified Eagle Medium (DMEM) supplemented with 10% bovine calf serum (BCS), 100 U/mL penicillin, 100 μg/mL streptomycin, and 2 mM L-glutamine. Cells were cultured at 37°C with 5% CO_2_ and ∼99% humidity. Viruses used in this study were described previously (12, 21).

### Cell culture infections

3T12 fibroblasts were infected with indicated viruses and MOIs. Briefly, viral stocks were diluted in low-volume (200 μL) of DMEM and added directly onto fibroblast cell monolayers seeded the previous day at a density of 2 × 10^5^ cells/well in 6-well plates. Plates were incubated at 37°C and rocked gently every 15 min for 1 hour. After 1hr of adsorption, cells were incubated in an appropriate culture volume of cDMEM for the indicated times at 37°C. For multistep growth curves, infected cells were frozen at −80°C at the indicated time points. Cells were subjected to freeze-thaw lysis to release progeny virions, and lysates were serially diluted for plaque assays as described (55).

### Plaque assays

Plaque assays were performed as previously described (55) using BHK21 cells. Briefly, infected cells were overlaid with 1.5% methylcellulose in DMEM supplemented with 2.5% calf serum, 100 U/mL penicillin, 100 μg/mL streptomycin, and 2 mM l-glutamine and incubated at 37°C for 4 to 6 days. Cell monolayers were stained with a solution of crystal violet in formalin for identification and quantification of plaques.

### CRISPR-Cas9 deletion of *Ifnar1*

Ifnar1 was deleted in 3T12 fibroblasts using the CRISPR-Cas9 system. The pLentiCRISPRV.2 (Addgene #52961) plasmid was digested with BsmBI-v2 and ligated with guide RNA sequences specific for Ifnar1 in **Table 3** (56). Cloned plasmids were transfected into HEK293T cells in combination with psPAX2 and pMD2.G plasmids using polyethyleneimine (Sigma). Supernatants were collected 72 hours post-transfection, passed through 0.45μm syringe filters and applied to 3T12 fibroblast in 6-well plates. 3T12 fibroblasts were transduced with retroviral supernatants by centrifugation for 30m at 1000 x *g* for 30m at 37°C in the presence of 10ug/mL polybrene (Santa Cruz Biotechnology). Cells were washed and media was replaced with cDMEM. 48 hours post-transduction, puromycin (Invitrogen; 2μg/mL) was added to the medium. Puromycin selected 3T12s were tested for Ifnar1 deletion by treatment with recombinant mouse IFN-β (Cell Signaling Technologies).

**Table 3.**
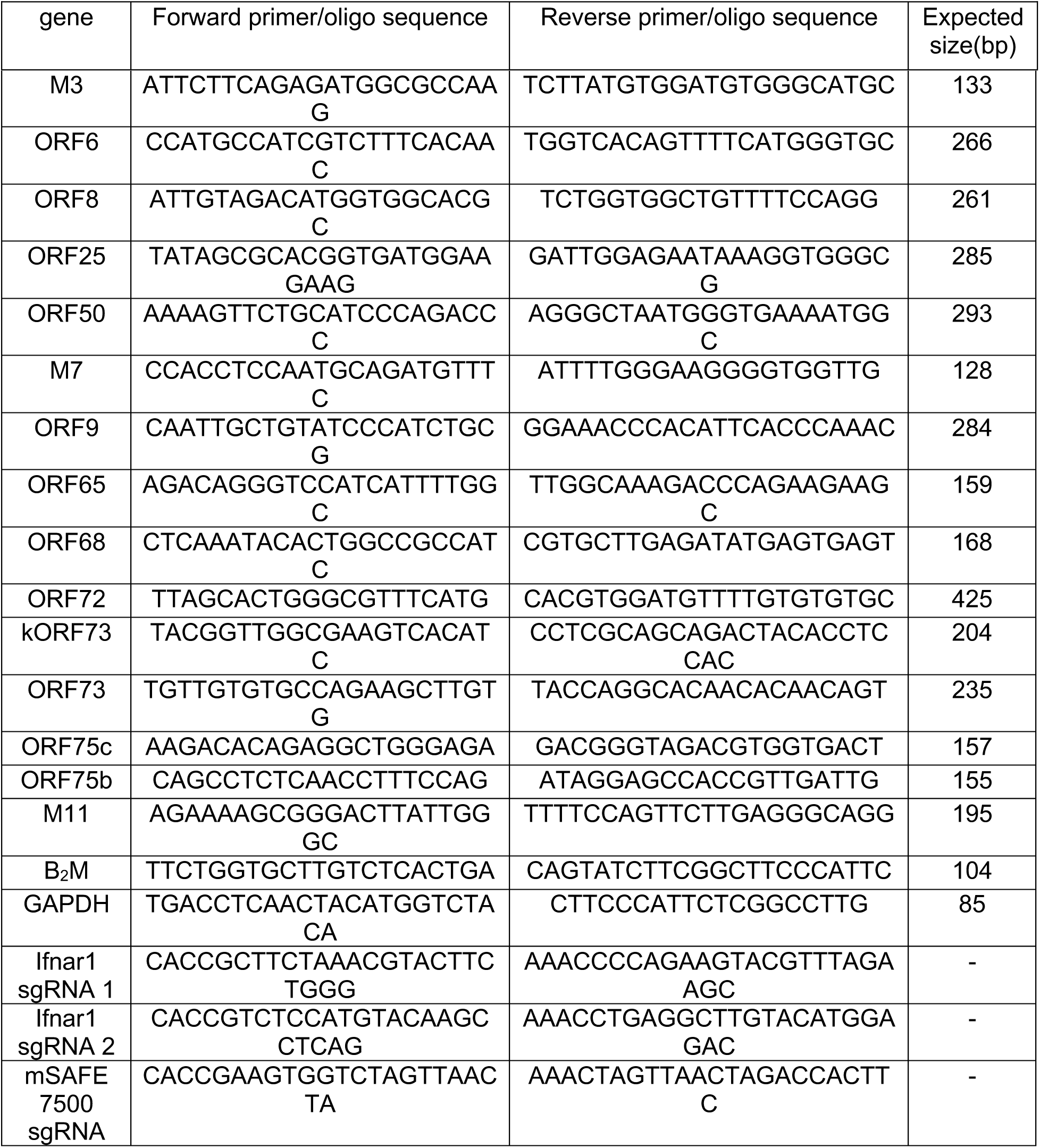
Primers for qRT-PCR and guide RNA oligonucleotide sequences for CRISPR/Cas9 genome editing.

### Splenocyte isolation and limiting-dilution assays

Spleens were homogenized through a sterile 40-μm filter and red blood cells were lysed by incubating tissue homogenate in 8.3 g/L ammonium chloride for 10 min at room temperature with shaking. Frequencies of cells harboring MHV68 genomes were determined using limiting-dilution (LD) PCR analysis with primers specific for an ORF50 locus as previously described (57). Frequencies of latently infected cells capable of reactivating were determined using a limiting-dilution analysis for cytopathic effect induced on an indicator monolayer as previously described (58).

### Immunofluorescence microscopy

3T12 fibroblast cells were plated infected as described above. Infected cells were washed with cold phosphate-buffered saline (PBS) and fixed in 10% phosphate-buffered formalin. Cells were permeabilized for 15 min with 0.5% Triton X-100–Tris-buffered saline (TBS) at room temperature. Supernatant was aspirated, and cells were incubated in blocking buffer (5% bovine serum albumin [BSA] and 0.1% Triton X-100, TBS) for 30 min at room temperature. Primary antibodies diluted in blocking buffer 1:1000 at 4°C overnight to probe for lytic antigens using serum from infected WT C57BL/6 mice. Samples were washed three times with cold wash buffer (0.1% Triton X-100, TBS). AlexaFluor 488-conjugated Goat anti-mouse secondary antibodies specific to primary antibodies diluted in blocking buffer 1:2500 and were added to the samples for 2 h in the dark at room temperature. Samples were washed three times with wash buffer and incubated with 1 μg/mL DAPI (4′,6-diamidino-2-phenylindole) (ThermoFisher) at room temperature to stain DNA. Cells were imaged by fluorescence microscopy using ×20 magnification on an Cytation 10 Confocal Imaging reader (Agilent).

### Immunoblot analyses

Immunoblot analyses were performed as previously described (59). Briefly, cells were lysed with radio immunoprecipitation (RIPA) buffer (150 mM NaCl, 20 mM Tris, 2 mM EDTA, 1% NP-40, 0.25% deoxycholate) supplemented with phosphatase and protease inhibitors. Samples were centrifuged at 16,000 × *g* to remove insoluble debris. The protein content for each sample was quantified using a DC protein assay (Bio-Rad). Samples were diluted in 4x Laemmli sample buffer and resolved by sodium dodecyl sulfate-polyacrylamide gel electrophoresis (SDS-PAGE) using 4-20% Mini-PROTEAN TGX precast gels (Bio-Rad) or 10% polyacrylamide gels prepared in-house. Resolved proteins were transferred to nitrocellulose membranes (Bio-Rad), blocked in either 5% BSA or 5% milk, and blots were probed with the indicated primary antibodies in **Table 3**. Antibody-bound proteins were recognized using horseradish peroxidase (HRP)-conjugated secondary antibodies (Jackson ImmunoResearch). Chemiluminescent signal was detected using a ChemiDoc MP imaging system (Bio-Rad) on blots treated with ProSignal Pico ECL reagent (Prometheus).

### RNA isolation and qRT-PCR

Total RNA was extracted from homogenized tissue or cells using a Quick-RNA Miniprep Kit (Zymo Research) according to manufacturer’s instructions. cDNA was synthesized using iScript cDNA sytheisis kit (Bio-Rad) according to manufacter’s instructions. For qRT-PCR, cDNA was diluted 1:3 and reverse transcription-PCR (RT-PCR) for *ORF50*, *ORF6*, *ORF9*, *ORF68*, *ORF25*, *ORF75c*, *ORF75b*, *ORF8*, *M7*, *M3*, *M11*, *ORF72*, *mORF73*, *kORF73, Actin*, and *B2m* transcripts was performed using SsoAdvanced Universal SYBR® Green Supermix (BioRad) using inner primers described in **Table 5** (60). Cycling conditions were 40 cycles of 94°C for 1 min, 62°C for 1 min, and 72°C for 1 min Applied Biosystems StepOnePlus PCR system.

**Table 4.**
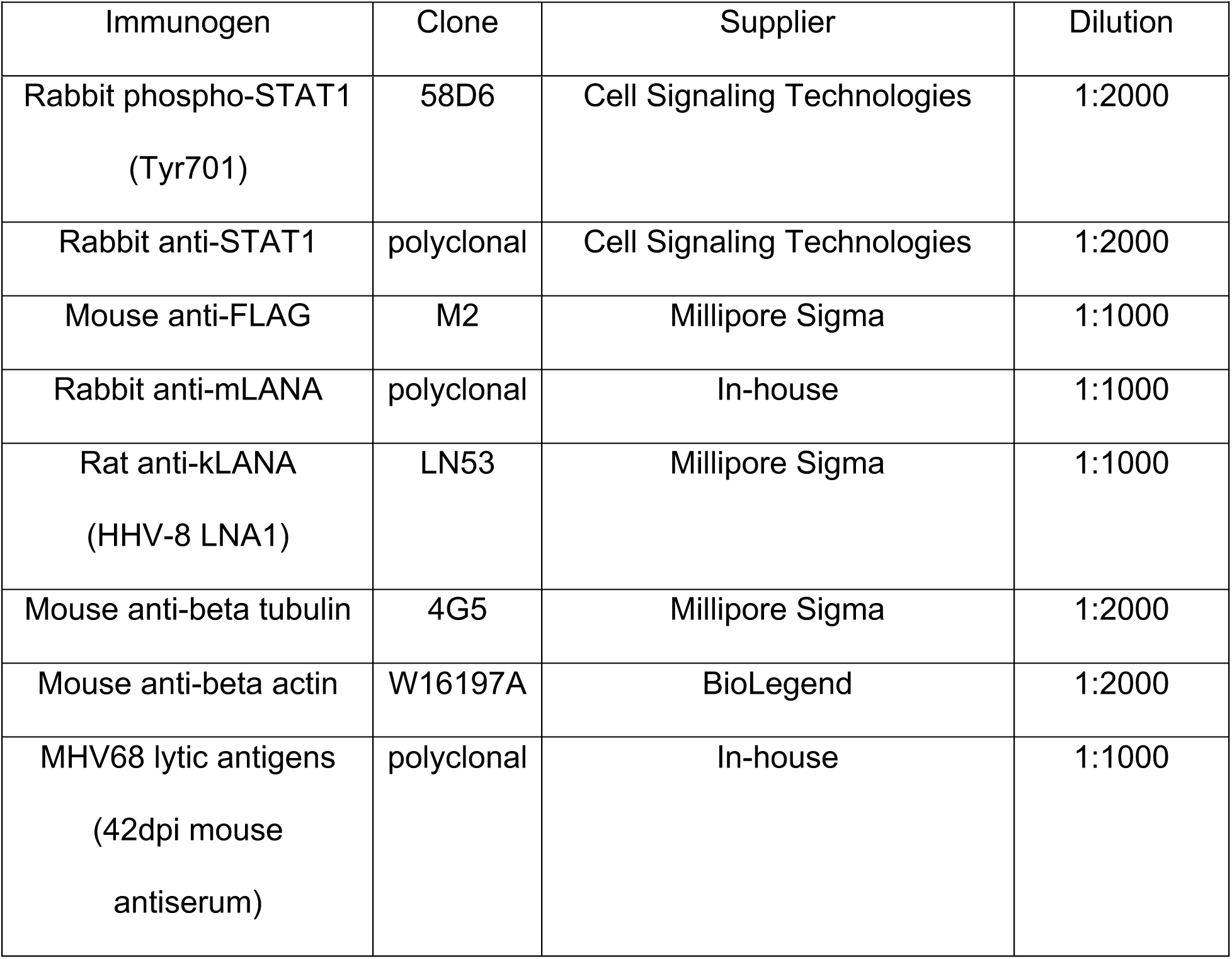
Antibodies used for immunoblot analyses.

**Table 5.** Antibodies used for flow cytometry analyses.

| **Table 5. Antibodies used for flow cytometry analyses.** |  |  |  |  |
| --- | --- | --- | --- | --- |
| Surface Marker | Clone | Fluorophore | Dilution | Supplier |
| anti-mouse<br>B220 | RA3-6B2 | rF710/PerCP-CY5.5 | 1:300 | TONBO |
| anti-mouse<br>CD19 | GD5 | BV650 | 1:300 | BioLegend |
| anti-mouse<br>GL7 | GL7 | APC | 1:300 | BioLegend |
| anti-mouse<br>CD38 | 90 | PE-Cy7 | 1:300 | BioLegend |
| anti-mouse<br>CD138 | 281-2 | PE | 1:400 | BioLegend |
| anti-mouse<br>CD3e | 145.2C11 | BV510/PerCP-Cy5.5 | 1:300 | TONBO |
| anti-mouse<br>CD11b | M1/70 | PerCP-Cy5.5 | 1:300 | TONBO |
| anti-mouse<br>CD11c | N418 | PerCP-Cy5.5 | 1:300 | TONBO |
| anti-mouse<br>TER-119 | TER-119 | PerCP-Cy5.5 | 1:300 | TONBO |
| anti-mouse<br>CD4 | RM4-5 | rF710 | 1:300 | TONBO |
| anti-mouse<br>CD8a | 2.43 | vF450 | 1:300 | TONBO |
| anti-mouse<br>CD44 | IM7 | PE, BV650 | 1:400 | BioLegend |
| anti-mouse<br>CD62L | MEL14 | BV605 | 1:300 | BioLegend |
| anti-mouse<br>PD-1 | J43 | PE-Cy7 | 1:300 | BioLegend |
| anti-mouse<br>CXCR5 | L138B7 | BV711 | 1:300 | BioLegend |
| anti-mouse<br>IFNAR1 | MAR1-5A3 | PE | 1:100 | BioLegend |

### Flow cytometry analysis and antibodies

3×10^6^ spleen cells were resuspended in 100 μl of FACS buffer (1X PBS, 2% BSA, 5mM EDTA) and blocked with anti-CD16/CD32 (BD Biosciences) for 15 minutes on ice. Cells were incubated on for 30 minutes with specific antibodies to identify surface markers. Lymphocyte subsets were identified with antibodies against indicated surface markers seen in **Table 5**. The data were collected on a BD LSR Fortessa or BD FACS Symphony flow cytometer (Beckman Coulter) and analyzed using FlowJoX v10.0.7 (Treestar Inc., Ashland, OR). Cells were first gated on lymphocytes based on forward and side scatter parameters and live cells were identified by exclusion of Fixable Viability dye eFlour™ 780 (eBiosceinces) prior to subgating.

### MHV68-specific tetramer staining

H-2D^b^-p56 MHC-peptide complexes were provided as biotinylated monomers by the NIH tetramer core facility and reconstituted with streptavidin-conjugated APC (used at a final dilution of 1:100). Tetramer staining was conducted in tandem with surface staining. Cells were fixed with 4% paraformaldehyde prior to data acquisition and analysis as described above.

### Enzyme-linked immunosorbent assays (ELISAs)

ELISA plates (MidSci) were coated with 0.5% paraformaldehyde-fixed viral antigens diluted 1:20 in carbonate buffer (0.0875 M Na2CO3, 0.0123 M HCO3, pH 9.2) and incubated overnight at 4 °C. Plates were washed in PBS with 0.05% Tween-20 (PBS-T) and blocked with 3% powdered milk-PBS-T for 1 hour at 37°C. Mouse sera was collected by cardiac puncture after euthanasia, and sera were serially diluted in 1% milk-PBS-T and incubated for 1 hour at 37°C. Virus-specific IgG was detected by incubation with HRP-conjugated donkey anti-mouse IgG (dilution 1:5000; cat# 715-035-150; Jackson ImmunoResearch). SureBlue substrate (KPL) was added for 3-5 minutes to detect virus-specific antibodies and read at 450 nm on a BioTek Synergy LX plate reader.

### Plaque-reduction neutralization assays (PRNTs)

Neutralization was tested by means of a plaque reduction as described (61). Briefly, sera were diluted 1:40 prior to three-fold serum dilutions in cDMEM. Each dilution was incubated with 25 PFU of MHV68 for 1 hour on ice. The virus/serum mixture was then added to a sub-confluent BHK21 monolayer (4 x 10^4^ cells/well) plated the previous day in a 24-well plate, in triplicate. Baseline consisted of virus standard alone. Infected cells were overlaid with methylcellulose and incubated at 37°C for 3-4 days. The overlay was removed, and the monolayers were stained with a solution of crystal violet (0.1%) in formalin to visualize plaques. Percent neutralization was determined by comparing of the number of plaques in experimental wells to no-serum controls.

### Statistical analysis

Data was analyzed using GraphPad Prism software (GraphPad Software, http://www.graphpad.com, La Jolla, CA). Total numbers of immune cells were analyzed by one-way ANOVA followed by post-tests depending on the parametric distribution. Based on Poisson distribution, frequencies of viral genome positive cells and reactivation were obtained from a nonlinear regression fit of the data where the regression line intersects 63.2%. Titer data were analyzed with an unpaired t-test.

### Ethics statement

Mouse experiments performed for this study were carried out in accordance with NIH, USDA, and University of Arkansas for Medical Sciences (UAMS) Division of Laboratory Animal Medicine and Institutional Animal Care and Use Committee (IACUC) guidelines. The protocol supporting this study was approved by the UAMS IACUC. Mice were anesthetized prior to inoculations and sacrificed humanely at the end of experiments.

## Acknowledgements

We thank Andrea Harris and the UAMS Flow Cytometry Core. This work was supported by R56AI150911 and R01AI181787 from the National Institutes of Health (NIH)-National Institute of Allergy and Infectious Diseases and start-up funds from the UAMS College of Medicine and Arkansas Biosciences Institute to J.C.F. S.J.M. was supported through a supplement to R01CA167065-S1 from the NIH-National Cancer Institute and R25GM083247 from the NIH-National Institute of General Medical Sciences (NIGMS). Flow Cytometry and Microscopy Cores and the work described here were also supported in part by the Center for Microbial Pathogenesis and Host Inflammatory Responses (P20GM103625) from NIH-NIGMS. The funders had no role in study design, data collection, and interpretation, or the decision to submit the work for publication.

S.J.M., S.M.O., and J.C.F. designed the study, analyzed the data, and wrote the manuscript. D.G.O designed and generated MHV68 BAC recombinants. J.C.F acquired the funding.

---

**Supplemental Figure 1.**
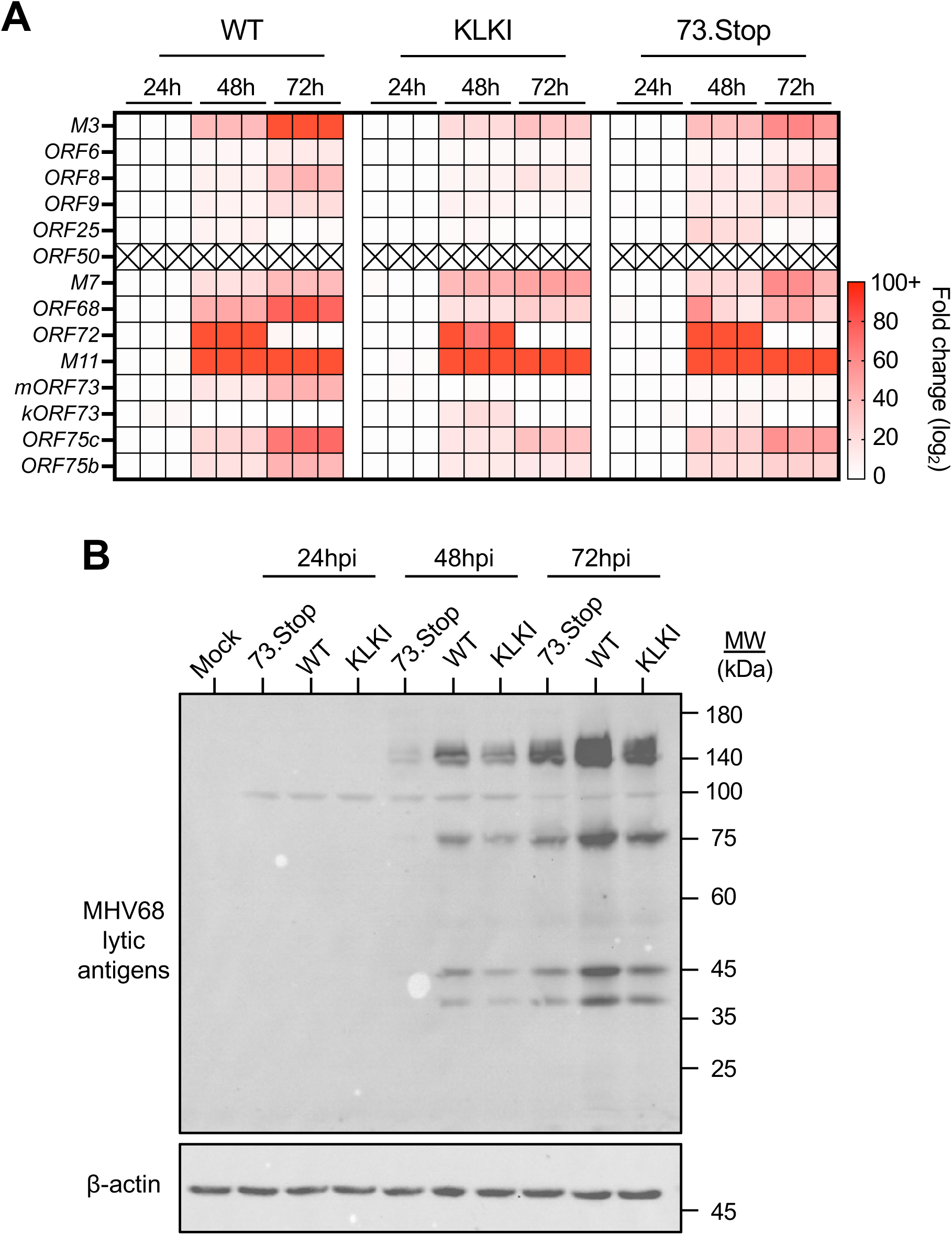
Expression of mRTA *in trans* rescues lytic antigen accumulation during KLKI MHV68 infection. RTA-expressing 3T12 fibroblasts were infected with the indicated virus at an MOI of 0.1 PFU/cell. (A) RNA was collected at the indicated times postinfection. Viral transcripts were quantified by qRT-PCR relative to *B2m* using the ΔΔCT method and normalized to transcript abundance at 24hpi. (B) Lysates were collected at the indicated times postinfection. Viral protiens were resolved by SDS-PAGE and immunoblot analyses were performed using antibodies that recognize the indicated proteins.

**Supplemental Figure 2.**
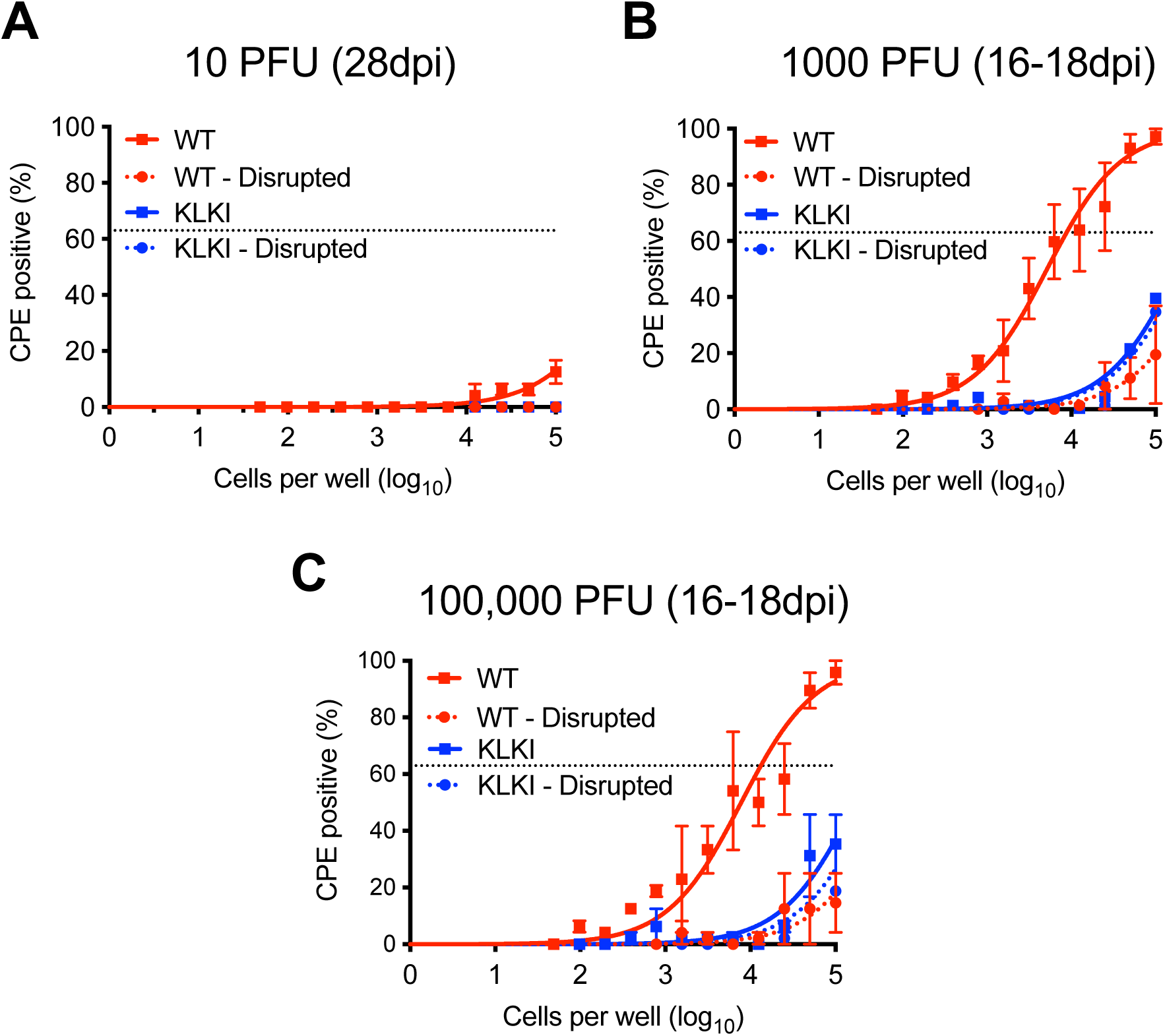
Reactivation from latency is reduced after KLKI MHV68 infection. C57BL/6 mice were infected IN with 10, 1000, or 100,0000 PFU of the indicated virus. Mice were sacrificed on day 16-18 postinfection (1000, or 100,000 PFU) or day 28 postnfection (10 PFU). Single-cell suspensions of splenocytes were serially diluted and reactivation frequencies were determined by *ex vivo* plating of splenocytes onto an indicator monolayer. Cytopathic effect was scored one-week postplating. Groups of 3 - 5 mice were used for each infection and pooled for analysis. Results are means of 2-3 independent infections.

**Supplemental Figure 3.**
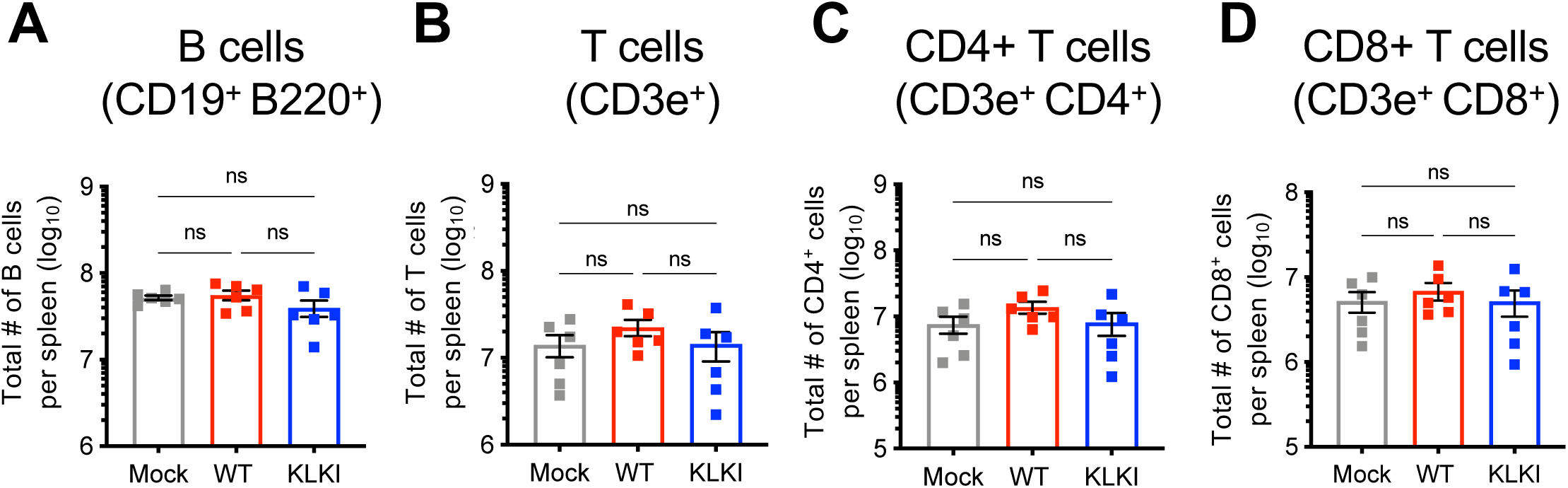
Total B and T cell populations are comparable between WT and KLKI MHV68 infection. C57BL/6 mice were mock-infected or infected IN with 1000 PFU of the indicated virus. Mice were sacrificed on day 18 postinfection. Total cells numbers were analyzed and quantified via flow cytometry. (A) B cells were defined as CD19^+^/B220^+^, (B) T cells as CD3e^+^, (C) CD4+ T cells as CD3e^+^/CD4+, and (D) CD8+ T cells as CD3e^+^/CD8^+^ cells. Groups of 3-5 mice were used for each infection and analysis. Results are means of 2 independent infections. Each dot represents one mouse. Error bars represent standard error of the mean. ns – not significant.

**Supplemental Figure 4.**
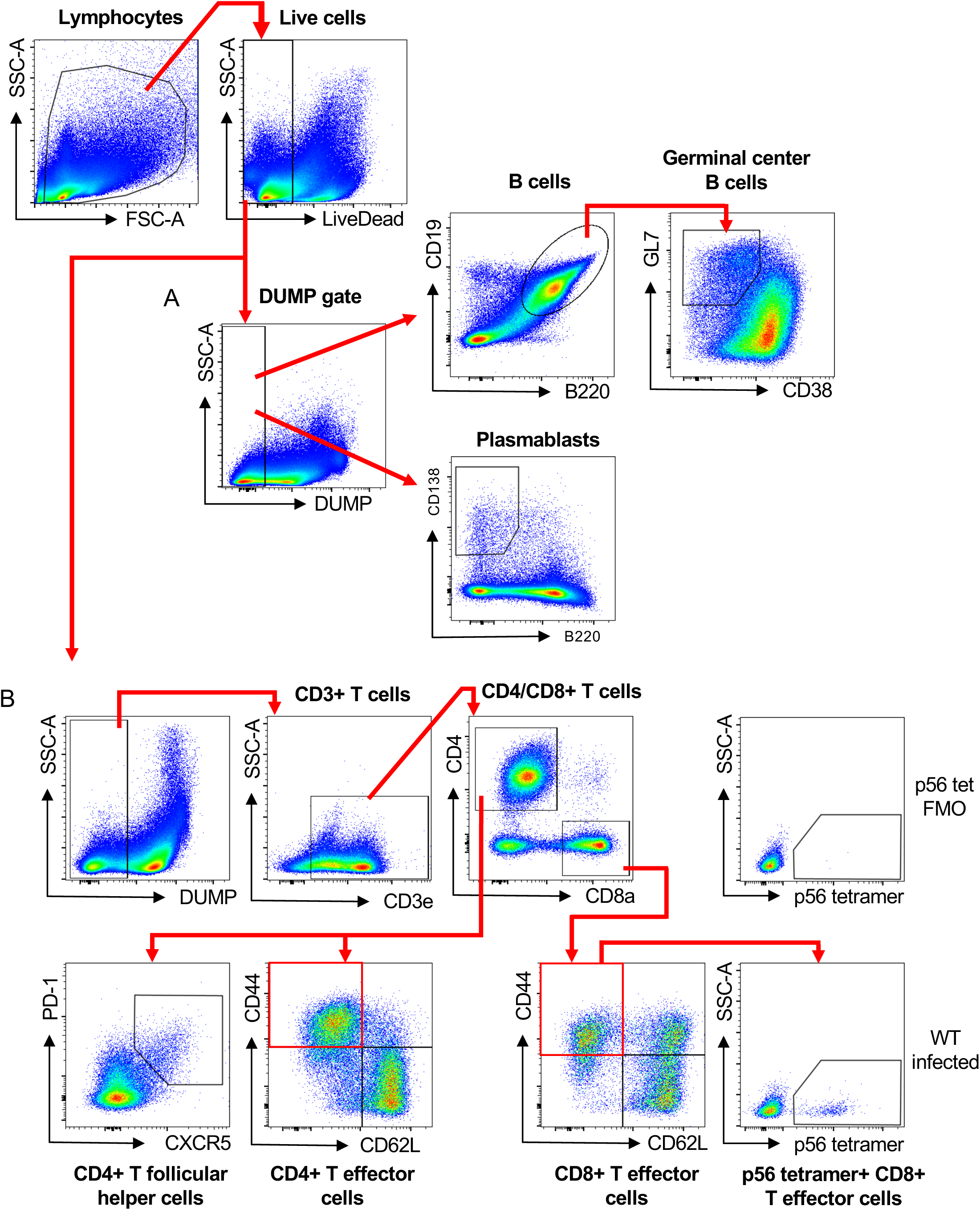
Flow cytometry gating strategies. C57BL/6 mice were mock-infected or infected IN with 1000 PFU of WT MHV68. Mice were sacrificed on day 16-18 postinfection and spleens were analyzed by flow cytometry. (A-B) Gating strategy for B and T cells populations. (A) For B cell populations DUMP exclusion gate was defined as viable CD3e^+^ CD11b^+^ CD11c^+^ TER119^+^ cells. (B) For T cell populations DUMP exclusion gate was defined as viable B220^+^ CD11b^+^ CD11c^+^ TER119^+^ cells.

